# A region-delineated snRNA-seq atlas of mouse spinal cord across lifespan resolves the interaction of normative aging programs with SOD1-G93A ALS

**DOI:** 10.64898/2026.04.08.717296

**Authors:** Michael Edison Pauli Ramos, Brijesh Kumar Singh, Oksana Shelest, Ian Tindel, Benisa Zogu, Ashley Dawson, Pranav Mathkar, Shaughn Bell, Ritchie Ho

**Affiliations:** Department of Biomedical Sciences, Cedars-Sinai Health Sciences University, Los Angeles, CA, USA; Board of Governors Regenerative Medicine Institute, Cedars-Sinai Health Sciences University, Los Angeles, CA, USA; Center for Neural Science and Medicine, Cedars-Sinai Health Sciences University, Los Angeles, CA, USA; Department of Neurology, Cedars-Sinai Health Sciences University, Los Angeles, CA, USA

## Abstract

Aging is the strongest risk factor for amyotrophic lateral sclerosis (ALS), yet how normative aging programs intersect with disease mechanisms remain unclear. Here we generated a lifespan-resolved, cell type-and region-specific single-nucleus RNA-sequencing atlas of the mouse spinal cord spanning embryonic development through advanced age in WT mice and end-stage disease in the SOD1-G93A ALS model. This resource enabled systematic comparison of physiological aging trajectories with disease-associated transcriptional changes across spinal cord cell types and rostrocaudal regions. We found that SOD1-G93A transcript and protein states differed markedly across spinal regions during disease onset and progression, and these molecular patterns paralleled the relative resilience of cervical regions and the heightened vulnerability of lumbar regions to degeneration in this transgenic mouse model. Prior to disease onset, we identified reduced ubiquitin expression that primed region-specific disruption of proteostasis in the SOD1-G93A spinal cord. Despite these disease-associated changes, aging-related transcriptional programs were largely preserved across most cell types, arguing against a global acceleration of aging in ALS. Instead, microglia emerged as a key exception, exhibiting accelerated and rewired aging- and disease-associated gene expression modules regulated by MITF and NRF2. Together, these findings provide an anatomically, cellularly, and temporally resolved framework for understanding how aging programs interact with disease-specific pathways to shape regional dysfunction and neurodegeneration in ALS.

## Introduction

Amyotrophic lateral sclerosis (ALS) is a late-onset neurodegenerative disease characterized by the progressive loss of motor neurons^1^. Although aging is the strongest epidemiological risk factor for ALS^2^ the relationship between physiological aging and disease pathogenesis remains incompletely understood. One possibility is that ALS reflects an acceleration of intrinsic aging programs, in which age-associated molecular processes are prematurely engaged or amplified^3^. Alternatively, ALS may arise through disease-specific mechanisms that intersect with, but are largely independent of, normative aging trajectories^4^. Discriminating between these models is essential for understanding disease initiation and progression. However, a temporally and finely resolved molecular roadmap of normative aging gene expression programs in central nervous system tissue is lacking.

Additionally, ALS exhibits pronounced anatomical and cellular heterogeneity. The site of motor symptom onset varies across patients and typically begins focally before spreading across the body^1^, reflecting differential vulnerability of motor neurons along the rostrocaudal axis and suggesting that spinal cord regions differ in their susceptibility to degeneration. Emerging evidence indicates that these differences may arise from cell-intrinsic transcriptional states and their interactions with local cellular environments^5^. However, it remains unclear whether such regional vulnerability is affected by baseline aging programs, disease-specific responses, or interactions between the two. Molecular hallmarks of aging including mitochondrial dysfunction, proteostasis decline, and altered stress responses are evident in ALS and may interact with disease-causing mutations to shape neurodegeneration^6,7^. Notably, many of these molecular changes precede overt phenotypic decline, highlighting the importance of defining early cell state transitions. However, human studies are inherently limited in resolving these dynamics, as ALS is typically diagnosed after symptom onset and longitudinal sampling of the prodromal phase is not feasible. Most transcriptomic studies in ALS patients rely on postmortem tissue, providing only endpoint snapshots and obscuring the temporal sequence of molecular events^8,9^. Experimental models therefore provide a critical framework for dissecting disease progression. The transgenic SOD1-G93A mouse model recapitulates key features of ALS, including stereotyped onset and regionally patterned motor decline, with hindlimb dysfunction preceding forelimb impairment^10,11^. These properties enable precise temporal and anatomical resolution of disease-associated changes and offer a unique opportunity to examine how molecular programs 1) evolve prior to and during neurodegeneration, 2) differ across the rostrocaudal axis of the spinal cord, and 3) intersect with age-associated changes.

Here, we generated a lifespan-resolved, cell type- and region-specific single-nucleus RNA-sequencing (snRNA-seq) atlas of the mouse spinal cord spanning embryonic development through advanced age in wild-type (WT) mice and end-stage disease in SOD1-G93A mice. This resource enabled direct comparison of age-and disease-associated transcriptional programs across rostrocaudal spinal cord cell types and regions. We defined when and where pathological processes diverge from physiological aging and identified cell type-specific deviations from healthy baseline gene expression patterns. Together, these analyses establish a temporally and anatomically resolved framework for distinguishing aging-associated from disease-specific molecular programs in ALS and for understanding how regional vulnerabilities contribute to selective neurodegeneration.

## Results

### A lifespan-resolved, cell type- and region-specific snRNA-seq atlas in WT and SOD1-G93A spinal cords

We first characterized disease progression in our cohort of SOD1-G93A mice. Weight gain began to slow at ∼P90, coinciding with the onset of motor decline measured by a modified Basso Mouse Scale^12^ (Fig. 1A). Consistent with prior reports, motor deficits were more pronounced in hindlimbs than forelimbs^11,13^ and indicative of region-specific vulnerability along the rostrocaudal axis. Prior studies have also reported the emergence of aging-associated markers at disease endpoint in these mice, suggesting accelerated engagement of cellular aging programs^14,15^. We thus hypothesized that mutant SOD1 in this mouse model accelerates cell type- and region-specific aging programs, leading to differential dysfunction and degeneration. To test this, we collected spinal cords from SOD1-G93A and age-matched WT littermates at presymptomatic (P25, P50), onset (P100), symptomatic (P125), and endpoint (P159) disease stages (Fig. 1B). To further define normative aging programs, we included a series of samples across WT animals at P200, P400, P600, and P800. All postnatal spinal cords (P25 to P800) were dissected into four anatomical regions that innervate distinct skeletal muscles along the body (Fig. 1B), and each segment was processed for snRNA-seq. We additionally profiled mouse induced pluripotent stem cells (iPSCs) and embryonic (E13.5) spinal cords to represent an embryonic developmental context for WT and SOD1-G93A samples.

**Figure 1.**
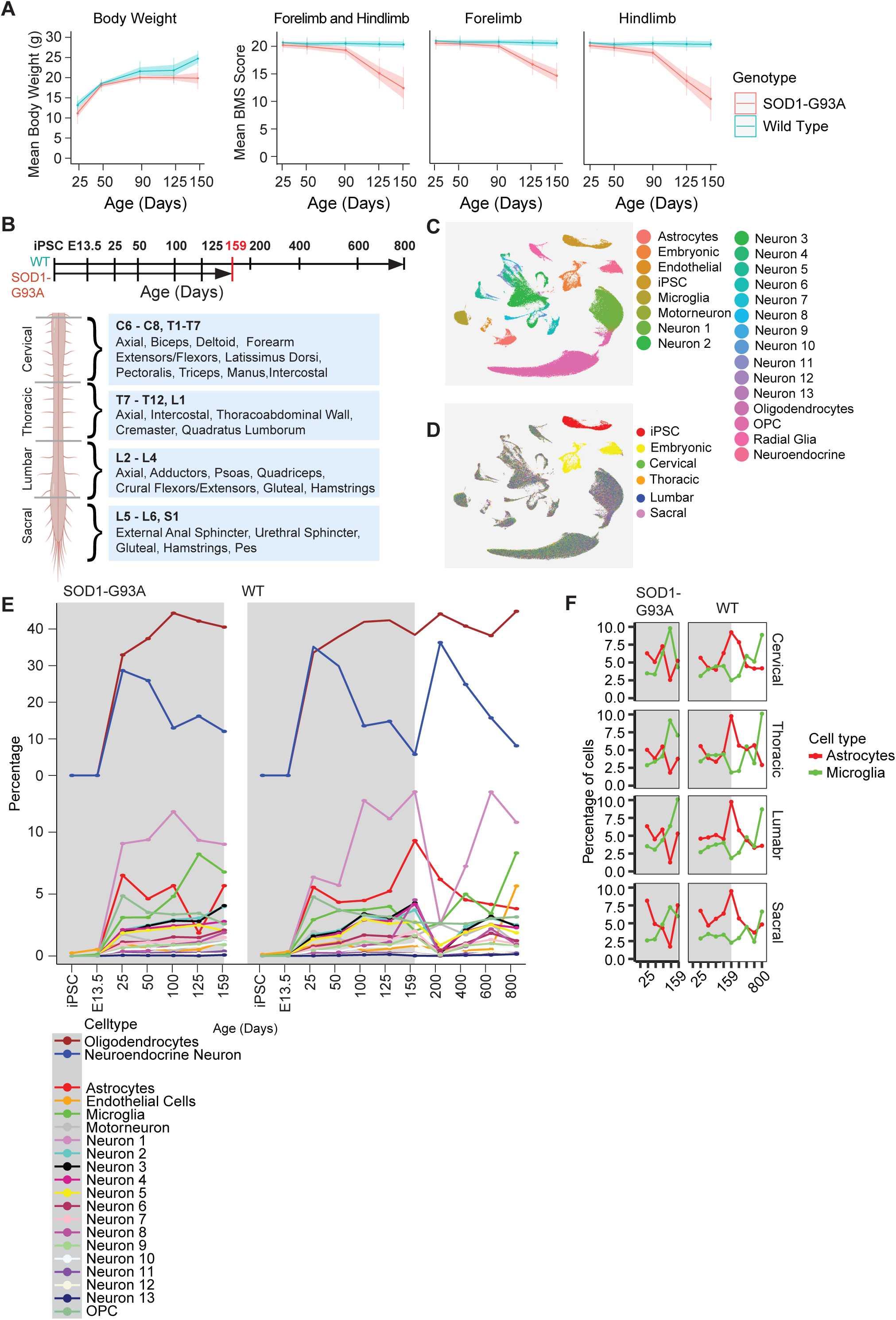
**Lifespan-resolved single-nucleus transcriptomic atlas of WT and SOD1-G93A spinal cord.** a, Body weight (left) and motor function trajectories for forelimbs and hindlimbs in WT (n = 5 mice per timepoint) and SOD1-G93A (n = 5 mice per timepoint) mice, showing onset of motor decline at ∼P90 and earlier hindlimb impairment. b, Experimental design and sampling strategy. Spinal cords were collected across presymptomatic (P25, P50), onset (P100), symptomatic (P125), and endpoint (P159) stages in SOD1-G93A mice, with extended aging timepoints in WT (P200–P800). Four anatomical regions were dissected along the rostrocaudal axis, each encompassing spinal segments and skeletal muscles they innervate. c, UMAP of 257,874 nuclei colored by annotated cell types, showing cell identity as the primary source of transcriptional variation. d, UMAP colored by spinal cord region, showing each postnatal cell type is evenly represented in each region. e, Cell type proportions across ages, showing overall stability across genotypes with preservation of major populations. Shaded grey regions represent ages spanning the disease course in transgenic mice. f, Astrocyte and microglia proportions across ages and regions, highlighting late-stage microglial expansion in SOD1-G93A mice.

On average, we sequenced ∼4,000 nuclei per sample. After filtering based on mitochondrial transcript proportion, unique molecular identifiers, and gene counts, we retained 257,874 high-quality nuclei annotated by genotype, age, and spinal region. We identified major cell types using canonical marker genes (Supplementary Fig. 1A). Two principal neuronal populations were observed: cells expressing high levels of *Nrg1*, *Nrg3*, *Rbfox1*, and *Rbfox3* were designated as “Neuron,” and cells enriched for *Syp*, *Snap25*, *Chga*, *Eno2*, and *Rtn1* were designated as “Neuroendocrine.” Harmonization with the spinal cord atlas from Sathyamurthy et al.^16^ revealed that Neurons aligned with multiple annotated neuronal subtypes, whereas Neuroendocrine cells clustered with populations previously labeled as “Unknown”. All other major cell types showed near-complete concordance with Sathyamurthy-annotated identities (Supplementary Fig. 1B). Motor neurons were identified by reclustering the Neuroendocrine population and isolating cells expressing *Chat* and

*Ache*, which formed a distinct cluster separated from the bulk of Neuroendocrine cells (Supplementary Fig. 1C). Consistent with these annotations, unsupervised clustering indicated that cell identity was the dominant source of transcriptional variation rather than genotype, age, and spinal cord region (Fig. 1C and 1D, Supplementary Fig. 1D and 1E). Cell type proportions were largely similar across SOD1-G93A and WT mice during early development and throughout the course of disease progression (P25 to P159), with the notable exception of microglia, which increased in abundance at later ALS stages (Fig. 1E, 1F, Supplementary Fig. 1F). This was consistent with heightened immune activation observed by others^17,18^. Altogether, this dataset captured transcriptional changes across lifespan and disease progression at cell-type resolution in all major spinal cord regions.

### SOD1-G93A transcript and protein states differ across spinal regions during disease progression

We first examined the expression dynamics of the *SOD1-G93A* transgene as well as endogenous mouse *Sod1* across all postnatal ages, spinal cord regions, and cell types within the SOD1-G93A genotype (Fig. 2A). *SOD1-G93A* transcripts were not detected in the nuclei extracted at presymptomatic ages (P25 and P50) but were induced starting at onset (P100) and were high throughout end stage (P159). Notably, induction of *SOD1-G93A* was highest in motor neurons and neuroendocrine neurons. These results suggest strong induction and sustained expression of the *SOD1-G93A* may impart higher levels of stress in neuronal cells or be a consequence of vulnerability. In the cervical region, expression of *SOD1-G93A* at P100 was chiefly lower than the subsequent symptomatic (P125) and end (P150) stages. In the thoracic, lumbar, and sacral regions, expression at P100 is chiefly higher than P125 and P150. This depicts a rostrocaudal effect on *SOD1-G93A* expression during paresis onset. In WT, endogenous *Sod1* was lowly expressed and largely unchanged throughout all corresponding regions and ages (Fig. 2A). Averaging nuclear *SOD1-G93A* expression across all cell types within each cervical and lumbar regions showed the same dynamic pattern (Fig. 2B). In contrast to the temporal expression patterns of RNA quantified from nuclei, image-based analysis of P50, P100, and P150 using RNAscope revealed that *SOD1-G93A* transcripts were already abundant at P50 and sustained through P150 in both regions, and they were predominantly cytoplasmic in spinal cord grey matter cells with relatively lower nuclear localization (Fig. 2C, Supplementary Fig. 2A, 2B). This suggests that at the presymptomatic stage, *SOD1-G93A* transcripts are efficiently exported from the nucleus and localize in the cytoplasm.

**Figure 2.**
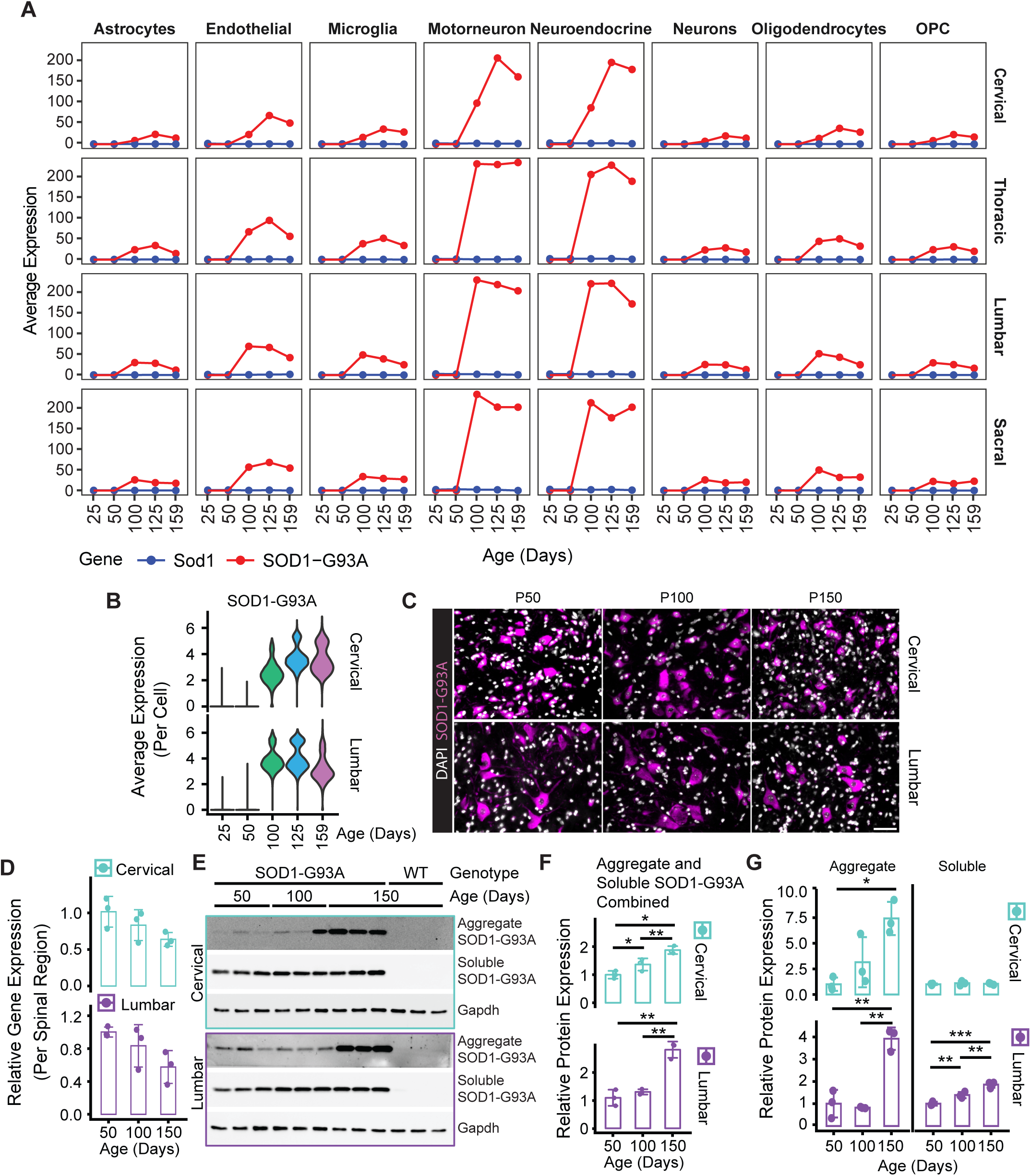
**Temporal and regional dynamics of SOD1-G93A expression and aggregation.** a, Expression of transgenic *SOD1-G93A* and endogenous Sod1 transcripts in the SOD1-G93A genotype spinal cord across disease time course (P25-159) and cell types, stratified by spinal region, showing highest expression in motor neurons and neuroendocrine cells. b, Violin plots of *SOD1-G93A* transcript levels across the disease time course in cervical (top) and lumbar (bottom) spinal cord, showing induction at disease onset (P100) and sustained expression thereafter, with earlier maximal expression in lumbar regions. c, RNAscope detection of *SOD1-G93A* transcripts in cervical (top) and lumbar (bottom) spinal cord, demonstrating predominantly cytoplasmic localization. Scale bars, 50 μm. d, Bulk qPCR quantification of *SOD1-G93A* transcript at P50, P100, and P150 (n = 3 mice per age), showing no significant change over time. Bars represent mean ± s.d.; points indicate biological replicates. e, Western blot analysis of insoluble (aggregated) and soluble SOD1-G93A protein in cervical and lumbar spinal cords from transgenic and WT mice at indicated ages (n = 3 mice per age). Soluble SOD1-G93A and Gapdh were imaged from the same blot; Aggregated SOD1-G93A was fractionated from the same sample protein extract but separately ran and imaged on another blot. SOD1-G93A was probed with an antibody specific to human SOD1-G93A mutant protein. f, Quantification of total SOD1-G93A protein in e (aggregate + soluble, normalized by Gapdh) over time, showing progressive accumulation in cervical and delayed increase in lumbar regions. g, Separate quantification of aggregated and soluble SOD1 in d normalized to Gapdh, showing continuous increase of soluble SOD1-G93A in lumbar spinal cord. Significance: *P < 0.05, **P < 0.01, ***P < 0.001, ****P < 0.0001.

However, this export efficiency is reduced at onset, symptomatic, and end stages (P100+), when nuclear *SOD1-G93A* transcripts accumulate and can be consequently measured by snRNA-seq. Likewise, this effect is stronger in the lumbar than cervical at P100. Quantitative measurement of overall *SOD1-G93A* levels in bulk spinal cord tissue using qPCR revealed a modest, non-significant decline in expression in both cervical and lumbar regions over disease progression (Fig. 2D). These data may reflect a progressive reduction of *SOD1-G93A* transcription combined with loss of *SOD1-G93A*-expressing neurons. In considering the temporal and regional patterns of snRNA-seq, RNAscope, and qPCR altogether, these observations suggest that the dynamic biogenesis and localization of *SOD1-G93A* is coincident with paresis onset and correlated with phenotypic severity.

Subsequent analysis of SOD1-G93A protein states during disease progression further revealed differential processing dynamics in cervical and lumbar regions. When aggregated and soluble fractions of protein extracts were analyzed together, total SOD1-G93A protein increased monotonically over time in the cervical region, whereas a significant increase in the lumbar region was observed only at endpoint (Fig. 2E, 2F). Separate analyses of aggregated and soluble forms of SOD1-G93A showed 1) a gradual increase of aggregated protein and static level of soluble protein in the cervical region and 2) a late-stage increase of aggregated protein and gradual increase of soluble protein in the lumbar region (Fig. 2G). We also noted the presence of aggregated SOD1-G93A in the lumbar spinal cord as early as P50 (Fig. 2E), suggesting that aggregation initiates earlier in this region and therefore begins from a higher baseline. By considering the differential timing of forelimb and hindlimb dysfunction, our observations support the concept that increased soluble and misfolded SOD1-G93A is deleterious, and aggregated SOD1-G93A is either benign or protective^19–21^. Overall, integrating transcript localization with protein state analyses revealed a temporally and regionally distinct regulation of SOD1-G93A biogenesis and proteostasis that paralleled the pattern of disease progression along the rostrocaudal axis of the spinal cord.

### Intrinsic reduction of ubiquitin primes region-specific proteostatic changes in SOD1-G93A spinal cords

The reliable timing of hindlimb paresis at ∼P90 in our SOD1-G93A mice enabled us to examine molecular changes occurring prior to onset. This could identify processes that may contribute to the differential protein states of SOD1-G93A in cervical and lumbar regions and explain their subsequent rates of forelimb and hindlimb motor dysfunction. To this end, we performed differential gene expression analysis between presymptomatic (P50) and onset (P100) stages for each cell type and region respectively within WT and SOD1-G93A spinal cords (Fig. 3A). Across cell types and regions, we observed a substantial overlap of differentially expressed genes (DEGs) across genotypes (Jaccard similarity = 0.476). These findings indicate that many transcriptional changes occurring between P50 and P100 reflect normal physiological processes associated with maturation in young adulthood, which may be amplified or differentially utilized in the context of SOD1-G93A conditions.

**Figure 3.**
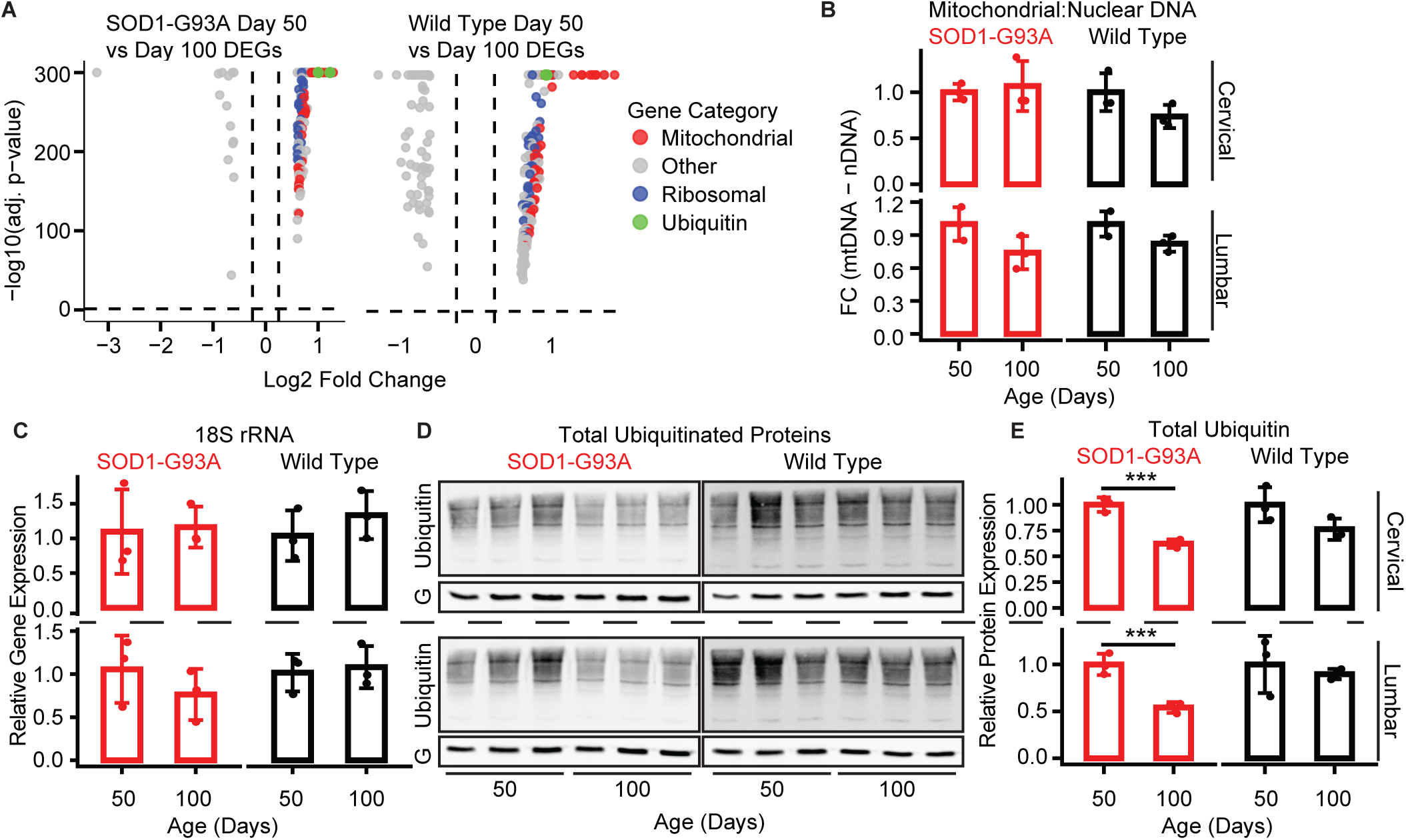
**Age-associated baseline decline in ubiquitin-dependent proteostasis may contribute to SOD1-G93A accumulation.** a, Volcano plots of DEGs between presymptomatic (P50) and onset (P100) stages in SOD1-G93A (left) and WT (right) spinal cord, showing significant genes corresponding to mitochondrial, ribosomal, and ubiquitin-related processes. b, Mitochondrial-to-nuclear DNA ratio in cervical and lumbar spinal cord, showing no significant change across the P50–P100 transition. c, *18S* rRNA levels as a proxy for ribosomal abundance, indicating preserved ribosome content despite reduced ribosomal gene expression. d, Western blot analysis of total ubiquitinated protein in cervical and lumbar spinal cord. G: Gapdh. e, Quantification of total ubiquitin levels normalized to Gapdh, showing a significant reduction in SOD1-G93A mice at P100 preceding detectable protein aggregation. Significance: *P < 0.05, **P < 0.01, ***P < 0.001, ****P < 0.0001.

Functional categorization of these shared DEGs revealed three predominant groups: genes associated with mitochondrial function, ribosomal biology, and ubiquitin-related processes (Fig. 3A). To determine whether the observed decline in mitochondrial gene expression at disease onset reflected a reduction in total mitochondrial content, we measured mitochondrial-to-nuclear DNA ratios. This analysis revealed a modest but non-significant decline in both cervical and lumbar spinal cord regions of WT and SOD1-G93A mice (Fig. 3B), suggesting that overall mitochondrial abundance is largely preserved despite reduced expression of mitochondrial components. Similarly, to assess whether decreased expression of ribosomal genes reflected a loss of total ribosome content, we measured 18S rRNA levels as a proxy for ribosomal abundance. No significant differences were observed between P50 and P100 in either genotype in both cervical and lumbar spinal cord regions (Fig. 3C), indicating that ribosome levels remain stable across this transition. Together, these findings suggest that reduced expression of mitochondrial and ribosomal genes may reflect a transient slowing or pause in biogenesis rather than loss of existing organelles or translational machinery.

In contrast, total ubiquitinated protein levels showed a significant decline in both cervical and lumbar spinal cord regions of SOD1-G93A mice, whereas WT mice exhibited only a modest, non-significant reduction (Fig. 3D and 3E). This indicates that despite reduced transcriptional output of ubiquitin at P100, WT cells are able to maintain this process of proteostasis. Since global protein ubiquitination is reduced in both SOD1-G93A cervical and lumbar spinal cords regions at P100, this highlights how these two regions handle SOD1-G93A protein differently; the cervical region shows a graded increase in aggregated but stable levels of soluble forms, while the lumbar region shows a latent increase in aggregated but graded increase in soluble forms (Fig. 2G). Thus, we intuit that intrinsic reduction of ubiquitin may prime SOD1-G93A spinal cords for dysregulated proteostasis.

### Cell type- and region-specific transcriptional responses in SOD1-G93A spinal cords

To examine the cell type- and region-specific effects of the mutant *SOD1* expression, we performed differential expression analysis between WT and SOD1-G93A cells at each prodromal and disease stage. DEGs segregated into two broad patterns: genes predominantly altered at endpoint, and genes differentially expressed at disease onset specifically in the cervical region (Fig. 4A, Supplementary Table 1). Because individual neuronal subtypes were low in abundance and yielded relatively few DEGs, all neuronal subtypes were pooled into a single group for subsequent analyses. Assessment of DEG overlap using the Jaccard similarity index revealed substantial sharing of both up- and down-regulated genes across major cell types (Supplementary Fig. 3A, 3B). This prompted us to examine whether these shared DEGs followed common temporal trajectories over time. Hierarchical clustering identified groups of genes with conserved temporal expression patterns across cell types (Supplementary Fig. 3C, Supplementary Table 2), suggesting the engagement of shared regulatory programs in response to disease. Motor neurons represented a notable exception; their limited numbers and progressive loss resulted in relatively few detectable DEGs. Nevertheless, the transcriptional changes we observed were consistent with previously reported SOD1-G93A-associated motor neuron signatures^22^ (Fig. 4B).

**Figure 4.**
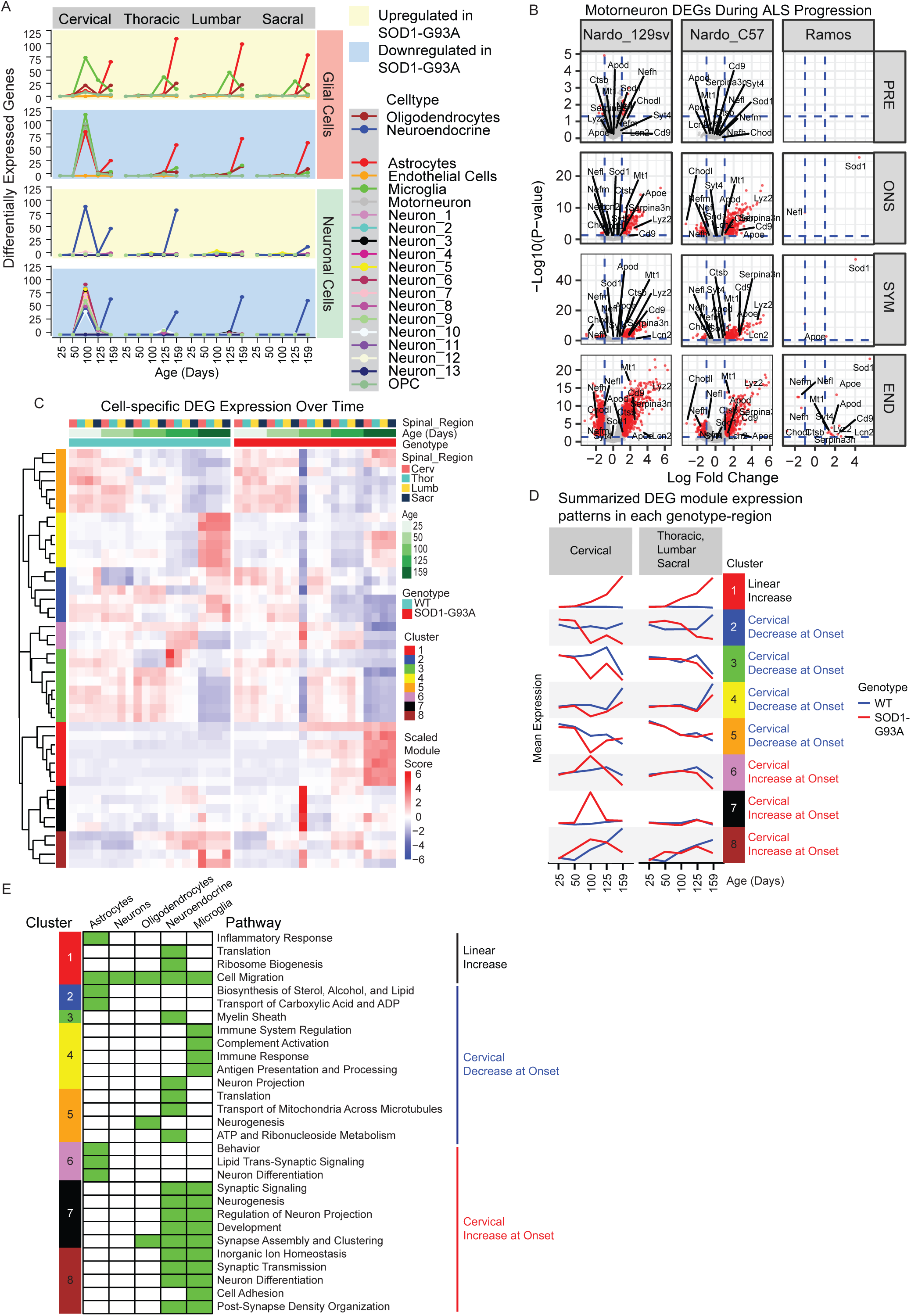
**Cell type- and region-specific transcriptional responses to SOD1-G93A.** a, Number of DEGs per disease stage and spinal region in SOD1-G93A compared to WT mice, showing predominant changes at endpoint and a distinct onset-specific response in the cervical spinal cord. b, Overlap of motor neuron DEGs with previously reported SOD1-G93A signatures in Nardo et al., 2013, confirming concordance with established disease-associated transcriptional changes. c, Hierarchical clustering of cell-specific DEG clusters across cell types and disease stages, revealing shared temporal expression patterns. d, Summary of DEG cluster expression with shared temporal dynamics across disease progression. e, Representative GO enrichment within cell type DEGs assigned to each cluster, highlighting which cells modulate specific pathways following the pattern of its corresponding cluster. See Supplementary Table 3 for a comprehensive list of enriched GO terms.

The shared temporal patterns across major cell types were consolidated into eight major DEG clusters (Fig. 4C, 4D), each enriched for distinct Gene Ontology (GO) pathways (Fig. 4E, Supplementary Table 3). Cluster 1, characterized by a monotonic increase in expression along all regions until end stage, shows enrichment for inflammatory response in astrocytes and cell migration across all cell types. This represents an overall tissue remodeling response associated with glial activation throughout disease progression. Clusters 2–5 exhibited a shared pattern of sharp downregulation specifically in the cervical region at P100 (paresis onset), while clusters 6–8 displayed transient upregulation specifically in the cervical region at disease onset. These changes did not occur or were delayed in the more vulnerable caudal regions. Downregulated pathways include sterol and lipid biosynthesis in astrocytes, immune response in microglia, and metabolically demanding activities in neuroendocrine neurons, while upregulated pathways include synapse formation and neurogenesis in neuroendocrine neurons, oligodendrocytes, and microglia. These findings suggest that region-specific engagement of shared cellular programs may contribute to differential neuronal resilience and help explain the delayed and less severe motor dysfunction observed in forelimbs compared to hindlimbs in SOD1-G93A mice.

### Aging-associated transcriptional programs are largely preserved in SOD1-G93A spinal cords

To determine whether aging processes are altered in SOD1-G93A mice, we first defined aging-associated transcriptional programs in WT cells from P25 to P800 and then compared them to those defined in SOD1-G93A cells from P25 to P159. We applied high-dimensional weighted gene co-expression network analysis (hdWGCNA)^23^ to each cell type and correlated module eigengene expression with age using a threshold (|r| ≥ 0.3) (Fig. 5A, Supplementary Table 4). In WT cells, we defined five modules positively correlated and eight modules negatively correlated with age, whereas in SOD1-G93A cells, we defined two modules positively correlated and one module negatively correlated with age. Among all cell types, microglia exhibited the strongest associations with age (|r| > 0.5) (Fig. 5A). Jaccard indices for intersecting each cell type-aging-associated module across genotypes indicated that most modules most were cell type- and genotype-specific (Fig. 5B). A notable exception was the microglial “blue” module, which showed substantial gene overlaps between WT and SOD1-G93A networks (Supplementary Fig. 4A).

**Figure 5.**
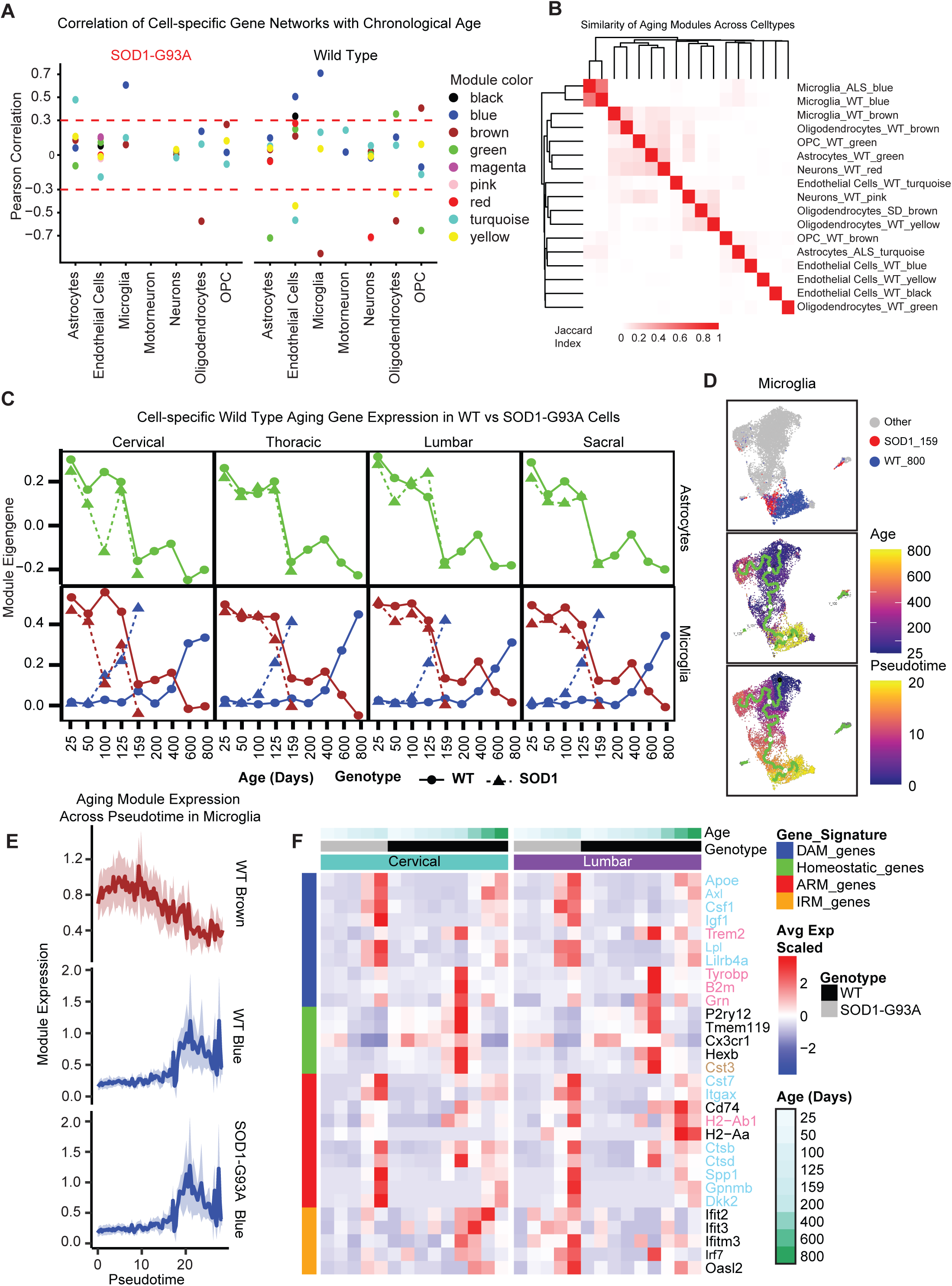
**Aging-associated transcriptional programs are largely preserved in SOD1-G93A mice with selective microglial divergence.** a, Correlation of module eigengenes with chronological age in SOD1-G93A (left) and WT (right) cells, identifying aging-associated gene networks. b, Jaccard similarity of aging-associated gene sets across cell types and genotypes, showing limited overlap except in microglia. c, Aging trajectories of WT-derived aging modules in WT (solid) and SOD1-G93A (dashed) cells, demonstrating preserved expression patterns of normative aging programs in astrocytes and accelerated aging in microglia. d, Microglia UMAP showing endpoint SOD1-G93A cells clustering with aged (P800) WT microglia (top), with corresponding chronological aging trajectory (middle) and pseudotime progression (bottom). e, Expression of the WT and SOD1-G93A microglial aging modules across pseudotime, showing accelerated engagement in SOD1-G93A microglia. f, Average expression of microglial disease-associated genes across the lifespan in cervical (left) and lumbar (right) regions. Font label colors indicate module membership; cyan: blue aging module in both WT and SOD1-G93A, pink: blue aging module in SOD1-G93A only, brown: brown aging module in WT.

In WT cells, negatively correlated aging modules identified across all cell types—except for weakly correlated modules in endothelial cells (r ≈ −0.3)—generally exhibited stable expression from P25 to P100, followed by a pronounced decline beginning at P125 (Supplementary Fig. 4B). This pattern indicates that transcriptional aging in the spinal cord is largely characterized by a coordinated reduction rather than activation of gene expression networks. Positively correlated aging modules in WT showed modest but sustained increases beginning at P125. Importantly, these age-associated gene network dynamics were highly consistent across cervical, thoracic, lumbar, and sacral regions, suggesting that the WT spinal cord ages uniformly across anatomical regions.

In SOD1-G93A cells, hdWGCNA identified three modules whose expression correlated with age: astrocyte turquoise (positive), microglia blue (positive), and oligodendrocyte brown (negative) (Fig.5A). The expression patterns for these modules tracked closely with chronological age (Supplementary Fig. 4B), and like the WT modules, their expression dynamics were consistent across spinal cord regions. Of these, only the microglial blue module showed substantial gene overlap with aging-associated modules identified in WT cells (Fig. 5B, Supplementary Fig. 4A), indicating that SOD1-G93A microglia engage and accelerate a conserved aging program relative to chronologically matched WT microglia (Supplementary Fig. 4B). GO analysis revealed enrichment of cell migration and cellular component movement (Supplementary Table 5). This illustrates microglia progressively adopt a migratory, injury-responsive state, likely reflecting active engagement with deteriorating motor neurons over the course of both WT aging and ALS. In contrast, SOD1-G93A astrocyte and oligodendrocyte modules in were distinct from those observed in WT (Fig. 5B), indicating these programs represent disease-specific responses rather than accelerated aging trajectories. GO terms enriched in the astrocyte module include cell motility, cell adhesion, extracellular matrix organization, and tissue development (Supplementary Table 5). This is aligned with the microglia response, and it also involved tissue remodeling. GO terms enriched in the oligodendrocyte module include cholesterol biosynthesis (Supplementary Table 5). This illustrates a progressive reduction in oligodendrocyte function in sterol and lipid production for myelination and metabolic support for motor neurons, this contributing to their vulnerability.

Using another means to assess whether SOD1-G93A cells enact WT aging programs, we calculated module eigengene scores for WT-defined aging modules in both genotypes. Across most cell types and regions, SOD1-G93A cells closely followed WT age-associated trajectories (Fig. 5C, Supplementary Fig. 4C). Interestingly in the cervical region, several WT aging modules, including astrocytes, oligodendrocytes, OPCs, neurons, and endothelial cells, were prematurely accelerated by P100 in the direction of their WT trajectory. However, they returned to their chronologically matched WT levels by P125. This distinct set of events may possibly confer some protective effects that could play a role in the resilience of forelimbs to motor dysfunction at this age. Microglia were an exception to this pattern; the WT blue module score increased more rapidly in SOD1-G93A microglia and monotonically increased until end stage in all regions (Fig. 5C), revealing a accelerated engagement of aging-associated transcriptional programs throughout the spinal cord. Overall, we conclude that the normative aging programs defined in WT spinal cords remain largely intact in SOD1-G93A spinal cords. This suggests that the *SOD1-G93A* transgene primarily drives neurodegenerative processes independent of widespread disruptions to normative age-related gene expression programs.

### Microglial aging- and disease-associated gene expression programs are accelerated in degenerating SOD1-G93A spinal cords

Because microglia were the only cell type exhibiting gene module divergence from WT aging module expression kinetics, we next examined their global gene expression pattern in greater detail. UMAP analysis revealed that endpoint SOD1-G93A microglia at P159 clustered with 800-day-old WT microglia (Fig. 5D).

Consistently, pseudotime trajectory analysis placed SOD1-G93A microglia further along the aging axis than chronologically matched WT microglia (Supplementary Fig. 4D). Module scores for microglial aging genes tracked closely with pseudotime progression, mirroring eigengene behavior across chronological age and supporting a role for aging module genes in driving global microglial state transitions (Fig. 5E, Supplementary Fig. 4E).

Both WT and SOD1-G93A microglial blue aging modules contained disease associate microglia (DAM) genes^24^ (*Apoe*, *Axl*, *Csf1r*, *Igf1*, *Lpl*, and *Lilrb4a*) as well as activated response microglia (ARM) genes^25^ (*Cst7*, *Itgax*, *Ctsb*, *Ctsd*, *Spp1*, *Gpnmb*, and *Dkk2*) (Fig. 5F). However, some DAM genes (*Trem2*, *Tyrobp*, *B2m*, and *Grn*) and one ARM gene (*H2-Ab1*) were uniquely assigned to the SOD1-G93A but not in the WT microglial blue aging module. Other ARM genes (*Cd74* and *H2-Aa*) were not assigned in either SOD1-G93A or WT microglial modules. In contrast, homeostatic microglia marker genes (*P2ry12*, *Tmem119*, *Cxcr1*, and *Hexb*) were not members of any microglial modules, except *Cst3* was assigned to the WT brown aging module. Interferon response microglia (IRM) genes^25^ were not assigned to any microglial modules. We then compared the lifespan expression pattern of these genes across genotypes and cervical and lumbar spinal cord regions (Fig. 5F). IRM genes (*Ifit2*, *Ifit3*, *Ifitm3*, *Irf7*, *Oasl2*) were selectively upregulated in the SOD1-G93A lumbar spinal cord as early as P50, prior to overt paresis. Expression of these genes progressively increased through P125 and reached peak levels by end-stage (P159). In WT lumbar spinal cords, these genes are moderately activated at P400 but the majority of these revert their expression by P600 and P800. In contrast, SOD1-G93A cervical spinal cord microglia exhibited low baseline IRM expression, with only a modest and transient increase at P125, returning toward pre-symptomatic levels by P159. There is notably strong expression of these IRM genes in WT cervical spinal cords at P200 that sustains throughout P600. These data indicate that a type I interferon–associated microglial program is initiated early and preferentially in the lumbar spinal cord, the region that undergoes earlier motor neuron degeneration in the SOD1-G93A mouse model. Notably, this interferon signature preceded widespread loss of homeostatic microglial identity.

Beginning at P100, coincident with symptomatic onset, SOD1-G93A microglia in both cervical and lumbar regions exhibited a progressive decline in homeostatic gene expression. This reduction continued through P125 and P159, indicating a broad destabilization of the surveillance microglial state during disease progression. The timing of this transition suggests that collapse of the homeostatic program represents a key inflection point in ALS pathogenesis and occurs independently of the early lumbar-restricted interferon response. At P125, SOD1-G93A microglia in both spinal regions upregulated DAM genes associated with lipid metabolism and phagocytic remodeling, including *Apoe*, *Lpl*, *Axl*, *Igf1*, *Csf1r*, and *Lilrb4a*. Expression of these genes further increased at P159, consistent with progressive engagement of a neurodegeneration-associated metabolic program. Strikingly, this transition occurred without induction of canonical Trem2-dependent DAM genes (*Trem2*, *Tyrobp*, *B2m*, and *Grn*). The absence of *Trem2* and *Tyrobp* upregulation indicates that the observed metabolic shift represents a Trem2-independent activation state distinct from classical amyloid-associated DAM described in models such as the 5xFAD mouse^24^. These data suggest that ALS spinal microglia engage a lipid-handling and trophic remodeling program that does not rely on transcriptional amplification of the Trem2 signaling axis. Concomitant with induction of metabolic genes at P125, we observed upregulation of ARM genes (*Cst7*, *Itgax*, *Ctsb*, *Ctsd*, *Spp1*, *Gpnmb*, and *Dkk2*). While ARM genes were modestly induced at P125, their expression increased further by P159, reaching levels comparable to DAM metabolic genes at end stage. ARM genes are associated with lysosomal protease activity, phagolysosomal remodeling, extracellular matrix signaling, and CD11c-associated activation^25,26^. Their progressive increase suggests escalating phagocytic load and lysosomal stress during late-stage disease. Importantly, ARM induction occurred in both cervical and lumbar regions, indicating that lysosomal remodeling represents a global late-stage feature of ALS spinal microglia rather than a regionally restricted phenomenon.

### SOD1-G93A rewires the regulatory interaction of SOD1, MITF, and NRF2 gene modules

Because WT aging and SOD1-G93A modules consist of co-expressed genes, we reasoned that they may be coordinated by shared transcriptional regulators. To investigate this, we constructed transcription factor regulons for each cell type using SCENIC^27^ and assessed their enrichment within aging modules identified in both WT and SOD1-G93A cells. To prioritize candidate regulators for experimental validation, we composed a weighted scoring framework that integrated multiple features indicative of functional relevance: (i) enrichment of a transcription factor’s predicted targets within a given aging module (weighted 0.8), (ii) correlation between regulon activity and module eigengene trajectories (0.3), (iii) expression level of the transcription factor in the corresponding cell type (0.6), and (iv) correlation between transcription factor expression and module eigengene dynamics across age (0.3) (Fig. 6A). Using this approach, we identified the top-scoring regulons for each cell type (Supplementary Fig. 5A) and characterized the biological pathways enriched among their target genes (Supplementary Fig. 5B; Supplementary Table 6).

**Figure 6.**
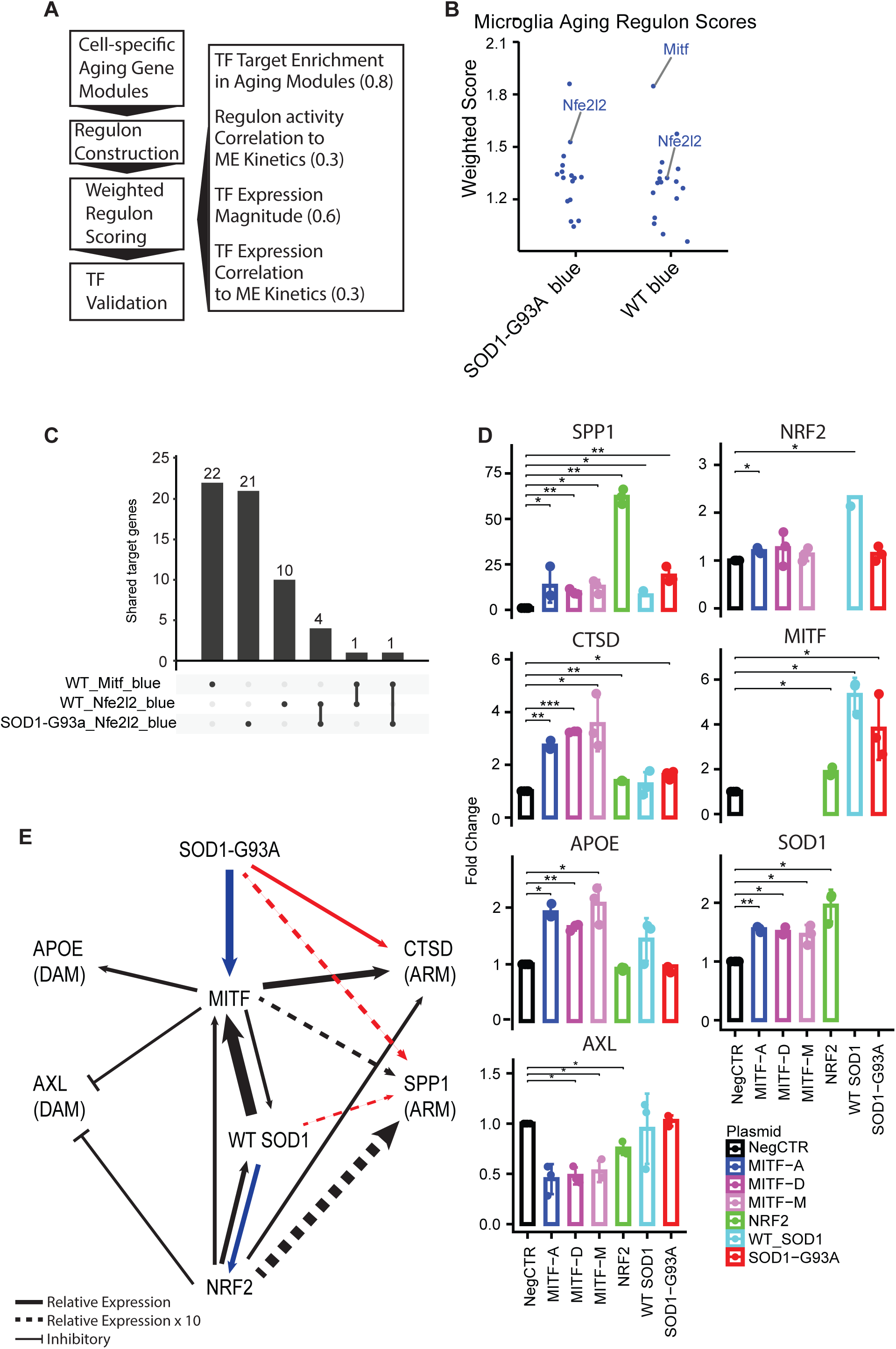
**Rewiring of MITF and NRF2 regulatory programs in SOD1-G93A microglia.** a, Schematic of weighted regulon scoring integrating target enrichment, expression, and correlation with aging module dynamics. b, Composite scores identifying top candidate transcription factors regulating WT and SOD1-G93A microglial aging modules, highlighting *Mitf* and *Nfe2l2* (Nrf2). c, Overlap of regulon targets with aging-associated genes in WT and SOD1-G93A microglia, showing divergence in regulatory programs despite shared module structure. d, Expression of representative DAM and ARM genes following overexpression of WT *SOD1*, *SOD1-G93A*, *MITF* isoforms, and *NRF2* in HEK293 cells, showing differential regulation of target genes as well as endogenous transcripts of each other. e, Regulatory model of *SOD1*, *MITF*, and *NRF2* in SOD1-G93A microglia, showing loss of coordinated stress-response regulation and selective activation of the ARM gene *CTSD*. Arrow thickness reflects fold-changes from panel d (solid: relative expression; dashed: relative expression x 10).

Given the pronounced disruption of aging trajectories in microglia, we focused subsequent validation efforts on this cell type. In the respective age-associated blue modules in WT and SOD1-G93A microglia, the highest-scoring regulons included *Nfe2l2* (Nrf2) and Mitf (Fig. 6B). Nrf2 (nuclear factor erythroid 2–related factor 2) is a master oxidative stress–responsive transcription factor that induces antioxidant, metabolic, and cytoprotective gene programs to maintain redox homeostasis^28^. In the SOD1-G93A mouse and other SOD1-linked ALS contexts, Nrf2 signaling is often dysregulated, influencing microglial inflammatory tone and their ability to buffer mutant SOD1–induced oxidative stress^29^. Mitf (microphthalmia-associated transcription factor) coordinates lysosomal biogenesis, lipid metabolism, autophagy, and phagocytosis in microglia and other immune cells^30^. It has also been identified as a critical driver of the DAM signature in neurodegenerative states^31^. We examined whether the predicted regulon target genes in each analysis overlapped with aging-associated genes within the microglial blue modules identified in WT and SOD1-G93A cells, as these modules are strongly correlated with age and enriched for disease-relevant signatures. Of the 429 aging genes in the SOD1-G93A microglial blue module, 26 were predicted targets of the Nrf2 regulon, representing a significant enrichment relative to the 4000-gene background (hypergeometric test, p = 3 × 10⁻¹²; OR = 6.4). Among the 408 aging genes in the WT microglial blue module, 24 were predicted targets of the Mitf regulon (hypergeometric test, p < 1 × 10⁻²⁰; OR ≈ 12.5), including disease-associated genes such as *Ctsd* and *Axl*.

Fifteen WT aging genes were targets of the Nrf2 regulon, which also showed significant but weaker enrichment (p = 2 × 10⁻⁴; OR = 2.8), indicating a shared but amplified Nrf2-associated program in SOD1-G93A microglia. Comparison of regulon targets revealed minimal overlap between WT and SOD1-G93A aging-associated genes (Fig. 6C). Taken together, these data demonstrate that while there is a high degree of overlap across the age-associated blue modules between WT and SOD1-G93A microglia, the respective gene regulatory activities of Mitf and Nrf2 are very different in each genotype.

Notably, Mitf regulon activity was detected within the aging-associated blue module only in WT microglia (Fig. 6B), suggesting that this component of the microglial aging program is absent or delayed in SOD1-G93A mice. In contrast, the Nrf2 regulon activity was detected within the respective aging-associated blue modules in both WT and SOD1-G93A microglia (Fig. 6B). However, Nrf2 regulated largely distinct gene sets between the two genotypes, with only four shared targets (*Fgr*, *Tank*, *Atf3*, and *Bcl2*), indicating divergent regulatory roles in normative aging versus disease conditions. Consistent with this distinction, Nrf2 regulon targets in WT microglia did not show significant GO enrichment, whereas Nrf2 targets in SOD1-G93A microglia were enriched for 94 GO terms, predominantly related to apoptosis, cell motility, tissue and muscle growth, and negative regulation of cell differentiation (Supplementary Fig. 5B, Supplementary Table 6). These data further support distinct gene regulatory activities between WT and SOD1-G93A microglia.

MITF has previously been shown to promote a disease-associated, highly phagocytic microglial state when overexpressed^32^, whereas NRF2 is a central regulator of antioxidant response element (ARE)–driven transcription and can act independently of SOD1 under oxidative stress conditions in immortalized cells^33^. Consistent with roles in aging-associated transcriptional regulation, both factors exhibited age-dependent changes in expression and regulon activity in microglia (Supplementary Fig. 5C–I). Notably, the activities of both the SOD1-G93A and WT Nrf2 regulons increased as each genotype reached the end of their lifespan (Supplementary Fig. 5F and 5G), consistent with increasing expression levels of *Nfe2l2* (Supplementary Fig. 5I). Interestingly, *Mitf* expression increased toward the end of lifespans for both SOD1-G93A and WT mice, but only the WT microglia revealed an increasing *Mitf* regulon activity, whereas SOD1-G93A microglia revealed no Mitf regulon (Supplementary Fig. 5H and 5I). These data again support a differential engagement of Nrf2- and Mitf-regulated pathways in SOD1-G93A conditions.

To further define how the SOD1-G93A mutation reshapes regulatory interactions among Nrf2, Mitf, and disease-associated gene modules, we examined the transcriptional relationships linking these factors to representative DAM (*Apoe*, *Axl*) and ARM (*Ctsd*, *Spp1*) genes that are upregulated in microglia of SOD1-G93A mice at disease end stage (Fig. 4C). To isolate cell-intrinsic transcriptional interactions, we used HEK293 cells to overexpress WT or mutant *SOD1*, *MITF* isoforms, and *NRF2*. Microglial models, including primary microglia, BV2 cell lines, or iPSC-derived microglia, exhibit strong baseline immune activation that can mask direct regulatory effects^34–36^. Using this reductionist system, we quantified how each factor could sufficiently influence the endogenous expression of the others, as well as representative DAM and ARM genes (Fig. 6D), and we summarized significant interactions (Fig. 6E). *MITF* overexpression significantly upregulated *APOE*, *CTSD*, *SPP1*, and *SOD1*, while suppressing *AXL* expression. *NRF2* overexpression significantly upregulated *CTSD*, *SPP1*, *MITF*, and *SOD1*, and similarly suppressed *AXL*, but did not induce *APOE*. These findings are consistent with *CTSD* representing a canonical MiT-family lysosomal target, whereas *APOE* and *SPP1* induction likely reflect indirect or secondary transcriptional effects rather than established direct MITF targets. The suppression of *AXL* by both *MITF* and *NRF2* further suggests antagonism between stress-adaptive lysosomal programs and interferon/injury receptor signaling modules.

WT *SOD1* overexpression significantly upregulated *SPP1*, *MITF*, and *NRF2*, revealing a reciprocal positive interaction between *SOD1* and these stress-adaptive transcription factors. In contrast, *SOD1-G93A* overexpression significantly upregulated *CTSD* and *SPP1* while attenuating *MITF* upregulation and failing *NRF2* upregulation. The direct comparison of WT *SOD1* and *SOD1-G93A* conditions revealed two major effects. First, WT *SOD1* positively and reciprocally reinforced *MITF* and *NRF2* expression, whereas this coupling was weakened (*MITF*) or lost (*NRF2*) with *SOD1-G93A*, indicating a partial loss-of-function in coordinated stress-responsive transcriptional regulation (Fig. 6E, relative magnitude of blue arrows). Second, *SOD1-G93A* conferred a gain-of-function effect characterized by enhanced induction of *CTSD* and amplified induction of *SPP1* (Fig. 6E, relative magnitude of red arrows), despite diminished *MITF* and *NRF2* support, indicating that their upregulation can occur independently of coordinated MiT-family or *NRF2*-driven programs. Taken together with the temporal kinetics of DAM and ARM gene expression observed in WT and SOD1-G93A microglia during aging and disease progression (Fig. 5F), these data support a model in which mutant SOD1 rewires regulatory interactions across MITF- and NRF2-centered modules. Specifically, SOD1-G93A appears to disrupt reciprocal reinforcement between MITF and NRF2 while promoting selective, stress-associated induction of ARM genes through alternative pathways, thereby contributing to the WT aging- uncoupled and disease-skewed activation state observed in ALS microglia.

## Discussion

Aging is the dominant risk factor for ALS, yet how age-associated molecular programs intersect with disease-specific pathology remains poorly resolved. Furthermore, ALS exhibits marked heterogeneity in the anatomical site of onset, suggesting that intrinsic differences between motor pools and their surrounding cellular environments shape vulnerability to neurodegeneration. With paresis stereotypically initiating in the hindlimbs between 90-100 days of age and progressing rostrally, the SOD1-G93A mouse model provides a uniquely tractable system to interrogate these interactions. This regional bias offers a powerful opportunity to probe why distinct spinal cord regions mount divergent molecular responses to the same pathogenetic insult. By generating a spinal region-resolved snRNA-seq atlas across the lifespan of WT and SOD1-G93A mice, we show that mutant SOD1 engages molecular programs that are both age-dependent and region-specific. We observed a progressive increase in nuclear SOD1-G93A transcript abundance over the course of disease; however, integration with RNAscope revealed that this increase reflects a shift in subcellular localization rather than transcriptional induction alone. At presymptomatic stages, SOD1-G93A transcripts are predominantly cytoplasmic, whereas at disease onset and progression they increasingly accumulate in the nucleus, which is detectable via snRNA-seq. This suggests possible burdens in normal function of nuclear machinery. Notably, this shift in transcript localization occurs earlier and more prominently in lumbar regions, consistent with their heightened vulnerability.

At the protein level, SOD1-G93A exhibits distinct regional dynamics in soluble and aggregated states. In the cervical spinal cord, total SOD1-G93A protein increases progressively over time, driven primarily by accumulation of aggregated species while soluble levels remain relatively stable. In contrast, the lumbar region shows a gradual increase in soluble SOD1-G93A and a pronounced late-stage accumulation of aggregated protein, despite evidence that aggregation is initiated early in this region. Given that paresis onset occurs near postnatal day 90 in this model, the presence of early aggregates in lumbar tissue, coupled with the delayed and region-specific accumulation of insoluble protein, suggests that aggregation alone is unlikely to be the primary driver of early dysfunction. Instead, our data supports a model in which soluble and misfolded SOD1-G93A species contribute more directly to toxicity, while aggregation represents a secondary, potentially compensatory process that sequesters these species. This interpretation is consistent with prior studies^21,37^ and is further supported by the distinct temporal dynamics observed across spinal regions. In particular, the lumbar spinal cord—corresponding to the site of earliest functional decline—exhibits early transcript relocalization and accumulation of soluble protein, whereas the cervical region shows a more gradual proteostatic response characterized by steady aggregate accumulation. Together, these findings highlight that both transcript handling and protein state transitions are spatially and temporally regulated, and that their interplay may underlie the rostrocaudal pattern of disease progression in SOD1-G93A mice.

Paresis onset also coincided with a decline in total ubiquitin availability, indicated through transcriptional and protein levels. Because ubiquitin supply is a key determinant of proteostatic capacity, this temporal ordering is consistent with the possibility that early limitations in ubiquitination constrain the efficient clearance of soluble misfolded SOD1-G93A, thereby predisposing cells to the later accumulation of insoluble aggregates. The presence of a similar, albeit weaker, age-associated decline in ubiquitin levels in WT mice suggests that diminishing ubiquitin availability may be a feature of normal spinal cord aging that is exacerbated in the context of mutant SOD1 expression. Consistent with this model, prior studies have shown that enhancing protein folding and ubiquitination capacity—such as through chaperone overexpression—improves motor neuron survival in the SOD1-G93A model^38^. Together with our observation of declining ubiquitin availability at disease onset, these findings support a model in which early SOD1-G93A–driven proteostatic stress, compounded by age-associated constraints on ubiquitin resources, precedes overt protein aggregation and contributes to cellular dysfunction. Whether such proteostatic limitations directly shape region-specific DEG patterns or arise downstream of distinct transcriptional programs remains an important question for future investigation.

Consistent with the region-specific proteostatic phenotypes, SOD1-G93A versus WT transcriptional responses were not uniform across the spinal cord. At disease onset, the cervical region exhibited a pronounced transcriptional response that was largely absent from thoracic, lumbar, and sacral regions. Given the delayed and attenuated forelimb paralysis characteristic of this model, this early cervical activation is consistent with the engagement of region-specific compensatory or adaptive programs that transiently buffer against neurodegeneration such as synapse formation and neuron differentiation. To address whether SOD1-G93A broadly accelerates WT aging programs, we used hdWGCNA to define aging-associated gene networks across spinal cord cell types in both genotypes. In WT mice, age-correlated modules across most cell types declined after postnatal day 125, indicating that spinal cord aging is dominated by the gradual loss of youthful transcriptional programs rather than the emergence of novel deleterious states. Importantly, these same aging-associated modules were largely preserved in SOD1-G93A cells at matched chronological ages, arguing against a global acceleration of intrinsic aging trajectories by mutant SOD1.

Microglia emerged as a striking exception to these patterns. We observed increased expression of IRM genes in SOD1-G93A microglia as early as P50 in the lumbar but not the cervical spinal cord, suggesting a presymptomatic role in hindlimb vulnerability to paresis. This was followed by decreased expression of homeostatic genes and subsequently increased expression of DAM and ARM genes in both regions. Whereas IRM and homeostatic genes respectively had region-specific and non-linear expression dynamics in WT aging microglia and were not co-expressed with gene modules, DAM and ARM genes ultimately increased expression in older WT microglia and were co-expressed with aging modules. Importantly, their activation occurs after hindlimb paresis onset (P100) and subsequent to the emergence of SOD1-G93A-driven molecular dysfunction and becomes prominent by P125. Thus, DAM and ARM gene activation appears to represent a reactive response that may subsequently amplify disease progression. These findings align with accumulating evidence that microglia integrate aging-related and disease-specific signals^39–41^ and, once engaged, can act as amplifiers of neurodegenerative pathology.

To identify candidate regulators underlying this accelerated microglial transition, we leveraged co-expression networks to prioritize transcription factors whose activity closely tracked aging-associated module dynamics. Functional testing revealed that mutant SOD1 perturbs the activity of NRF2, a master regulator of antioxidant stress responses and thereby redox buffering capacity. This observation is consistent with prior reports demonstrating that NRF2 activation can become uncoupled from oxidative stress signaling in disease contexts^33^. While most cell types largely preserve WT-like aging trajectories, SOD1-G93A microglia exhibited both temporal acceleration and divergent gene regulatory activities, engaging aging-associated modules earlier and progressing more rapidly towards late-life transcriptional states. At the regulatory level, this divergence is seen in the decoupling of MITF- and NRF2-centered transcriptional networks. The loss of MITF and NRF2 activity, combined with gain-of-function induction of stress-responsive genes independent of these regulons, provides a mechanistic framework for how microglia transition into a disease-skewed state despite retaining some elements of the underlying aging program. These observations suggest that SOD1-G93A-driven ALS selectively co-opts pre-existing aging pathways, amplifies but also redirects them, ultimately promoting degeneration.

Several limitations should be considered. First, our transcriptomic profiling relied on dense temporal sampling with one mouse per timepoint, prioritizing resolution of dynamic state transitions over within-timepoint replication. While this design limits power to assess inter-individual variability, the highly stereotyped progression of the SOD1-G93A model and orthogonal validation approaches mitigate this concern. Second, regulatory directionality was assessed using overexpression assays in a heterologous HEK cell system, which does not fully recapitulate microglial identity. These experiments were intended to assess sufficiency rather than physiological relevance, and future *in vivo* or microglia-specific perturbations will be required to refine these regulatory relationships in a cell type-specific context. Finally, while the SOD1-G93A model provides a robust platform for studying mutant SOD1–driven pathology, its aggressively shortened lifespan does not fully capture the complex aging milieu of human ALS. The relative preservation of intrinsic aging trajectories observed here suggests that this model may underrepresent interactions between physiological aging and disease mechanisms relevant to sporadic ALS, underscoring the need for complementary systems. Future studies applying our framework in other transgenic mutant SOD1 mice with more delayed onset of motor dysfunction, such as the low copy number SOD1-G93A or SOD1G37R mice^42,43^, will help further define the relationship between aging and pathogenic ALS mechanisms.

Taken together, our findings support a model in which physiological aging and mutant SOD1–driven pathology largely proceed in parallel within the spinal cord, intersecting selectively in specific cellular contexts rather than through a global acceleration of aging. Microglia emerge as a critical point of convergence, where age-associated vulnerability, impaired redox signaling, and proteostatic stress synergize to drive maladaptive immune activation. These insights suggest that therapeutic strategies aimed at broadly “slowing aging” may be insufficient. Instead, targeting early, cell type-specific vulnerabilities, particularly those governing proteostasis, oxidative stress responses, and immune activation in microglia, may offer a more precise approach to delay disease onset or progression. By resolving aging- and disease-associated programs at single-cell resolution across the lifespan, this work provides a conceptual framework for disentangling age-related risk from disease-specific mechanisms in ALS and other late-onset neurodegenerative disorders.

## Acknowledgements

We thank the Cedars-Sinai Genomics Core for sequencing support. RH was supported by grants from the National Institute on Aging (R00AG056678), Center for Research in Women’s Health Sciences Research Award by the Louis B. Mayer Foundation, and Winnick Foundation Research Scholars Award. MEPR and BKS were supported by the California Institute for Regenerative Medicine (EDUC4-12751).

## Author Contributions

RH and MEPR conceptualized and designed the study. MEPR, BKS, OS, BZ, AD, PM, SB, and RH performed experiments. OS conducted behavioral testing, and IT analyzed behavioral data. MEPR carried out data processing and analysis. RH supervised all aspects of the study. RH and MEPR wrote the manuscript, with editing input from BKS.

## Ethics Approval

All animal procedures were approved by the Institutional Animal Care and Use Committee (IACUC) at Cedars-Sinai Medical Center under protocol number 7496.

## Sex as a Biological Variable

Female mice were used exclusively to match the sex of previously published spinal cord microarray data sets, enabling direct comparison of transcriptional profiles and reducing variability from sex-dependent differences.

## Methods

### Animals

All procedures were approved by the Cedars-Sinai Institutional Animal Care and Use Committee (IACUC protocol 7496). Heterozygous B6.Cg-Tg(SOD1*G93A)1Gur/J transgenic male mice^10^ were crossed with WT C57BL/6 females, with pairings maintained at least five cousins apart. Mice were group-housed in ventilated cages under a 12-hour light/dark cycle with ad libitum access to standard chow and water.

Environmental enrichment (e.g., nesting material, shelters) was provided. Genotyping for the SOD1-G93A transgene and sex determination were performed by Transnetyx using standard probe sets. Female littermates were used for pairwise comparisons between WT and transgenic mice. Older WT mice for late timepoints (p400 and above) were purchased from The Jackson Laboratory and aged in-house. No a priori sample size calculations were performed. Mice that failed neurological scoring criteria were excluded. Assessors were aware of animal sex, age, and genotype during testing. For snRNA-seq, one female mouse was used per genotype and timepoint; each spinal cord was sectioned into cervical, thoracic, lumbar, and sacral regions and processed independently. All subsequent molecular experiments were performed using female mice. RNA, DNA, and protein analyses were conducted using three independent female biological replicates per condition.

### Phenotypic assessment of SOD1-G93A mice

Body weight was monitored longitudinally as a prognostic indicator of disease onset. Motor function was evaluated using a modified Basso Mouse Scale (BMS) analysis^12^. Assessments were performed at 25, 50, 90, and 125 days of age, and at humane endpoint (≥150 days) for SOD1-G93A mice. Humane endpoint was defined by loss of righting reflex (inability to right within 30 seconds when placed on the side) or paralysis resulting in impaired feeding, drinking, or ambulation. Animals that failed neurological scoring criteria were excluded from analysis. Assessments were performed in the home cage environment on bedding without obstacles. Locomotor function was assessed using the BMS during a 3-minute open-field task. To increase sensitivity to gradual disease progression, the original ordinal BMS was converted to a ratio-based scale, enabling independent scoring of limb movement, weight support, stepping, coordination, trunk stability, and tail position. Scores were averaged across limbs for each animal.

### Spinal Cord Dissection and Nuclei Isolation

Mice were sacrificed at assigned timepoints. Mice were put to sleep by dousing gauze in isoflurane in an oscillating vial placed on the mouse snout. The mouse is then perfused with PBS until the internal organs become pale. The spinal cord is extracted using hydraulic extrusion. Mouse spinal cords were flash frozen in dry ice and stored in −80 °C. Time from isolation to freezing was minimized as much as possible, with the spinal cords immediately going on dry ice after extrusion. Gross anatomical sectioning was conducted by cutting at the lumbar enlargement, then further cutting the two pieces in half on dry ice. Samples are defined as nuclei isolated from a specific region (cervical, thoracic, lumbar, sacral) of a certain genotype at a specific age (e.g. WT_p100_Cerv). Each sample is processed independently and represents one biological replicate per age and genotype, with regional dissections treated as technical subdivisions of the same animal. Nuclei isolation was performed on each sample as previously described with minor modifications^44^. Tissues were dounced in 2 ml of homogenization buffer (0.016 mM PMSF, 0.167 mM beta-mercaptoethanol, 320 mM Sucrose, 0.1 mM EDTA, 0.1% NP40 (Millipore Sigma, Cat# 18896-50 ml), 40 units per sample Roche RNase Inhibitor (Millipore Sigma, Cat# 3335399001), 1 tablet Roche Protease inhibitor (Millipore Sigma, Cat# 4693116991)) 50 times using the thinner pestle and another 50 times with the thicker pestle. The homogenate supernatant is then filtered through a 70-micron porous Flowmi filter (Millipore Sigma, Cat# BAH136800070-50EA). This is then gently mixed with 50% iodixanol solution to make a final solution of nuclei suspended in 25% iodixanol (25% Opti-prep (Millipore Sigma, Cat# D1556-250 ml) in homogenization buffer) for gradient isolation of nuclei. This is then added on top of iodixanol solution layers containing 40% iodixanol containing Orange G (40% opti-prep in homogenization buffer, 160 mM Sucrose) for visual clarity at the bottom, 29% iodixanol (29% Opti-prep in homogenization buffer) in the middle, and 25% iodixanol containing nuclei at the top. We centrifuged this for 1 hour at 3000 g at 4 °C with no brakes then collected the nuclei from the interphase between the 40% and 29% iodixanol layers. These were diluted in 1% BSA solution containing RNAse Inhibitor and centrifuged for 10 minutes at 500 g at 4 °C. The nuclei are then counted using trypan blue and hemocytometer.

### Library Preparation and Sequencing

snRNA-seq libraries were prepared using the Chromium Next GEM Single Cell 3’ Reagent Kits v3.1 (10x Genomics) according to the manufacturer’s protocol. Nuclei suspensions were loaded onto the Chromium controller aiming to recover approximately 4,000 nuclei per sample. Multiplexing was performed using the Chromium i7 Single Index Kit, and resulting cDNA libraries were quantified using an Agilent Bioanalyzer. Libraries were sequenced on an Illumina NovaSeq 6000 platform using paired end reads (28 bp Read 1, 8 bp i7 Index, and 91 bp Read 2) to a depth of approximately 50,000 reads per nucleus.

### Data Analysis Preprocessing

Raw sequencing data were processed using Cell Ranger v5.0.1 (10x Genomics). Sample demultiplexing was performed using the sample-specific i7 index barcodes. The cellranger count pipeline was run using the --include-introns flag to capture unspliced nuclear transcripts, which are typical in snRNA-seq data. Reads were aligned to the mouse genome (mm10) using the 10x Genomics reference refdata-gex-mm10-2020-A. The --expect-cells=4000 parameter was used to guide cell calling. For each sample, the filtered gene-barcode matrices output by Cell Ranger were used for downstream quality control, normalization, and analysis in R using Seurat (v4.3.0)^45^. Single cells were identified by filtering for quality using mitochondrial transcript percentage, total UMI count, and total number of genes detected per cell. Each of these metrics was transformed into a Z-score within each sample to control for sample-specific variability in sequencing depth and complexity. Specifically, mitochondrial transcript percentage (percent.mt), number of genes (nFeature_RNA), and total transcript counts (nCount_RNA) were computed using Seurat. Z-scores were calculated per sample using the base R scale() function, and added to the Seurat object as metadata. Cells were retained only if they had Z-scores between –3 and +3 for all three metrics. This approach ensured consistent filtering across samples while removing low-quality or outlier cells (e.g., apoptotic cells, doublets, or low-complexity transcriptomes). Following QC, gene expression data were normalized using Seurat’s NormalizeData() function. The top 2,000 most variable features were identified using FindVariableFeatures(), and the data were scaled using ScaleData(). Principal component analysis (PCA) was then performed using the RunPCA() function. An elbow plot was used to determine the number of principal components (PCs) to retain, with downstream analyses typically using the top 12 PCs based on visual inspection. Cell clustering was performed as described in Seurat package version 4.3.0. The default number of variable genes, 2000, was used. Clustering resolution was reduced until main cell clusters were visibly identifiable in the UMAPs generated using Seurat’s RunUMAP() function.

### Cell type identification

The FindAllMarkers() Seurat command was used to identify cell-specific markers expressed by cell clusters. Cell types were identified through expression of canonical cell markers. A large population of cells that expressed neuronal markers but were entirely separate were labeled “Neuroendocrine Neurons” due to expression of neuroendocrine genes *Rtn1* and *Nse*. These cells clustered very closely to motor neurons but did not express cholinergic markers *ChAT* or *Ache*. Because accurate cell type identity was critical, cells lacking expression of their assigned cluster’s marker genes were excluded from the dataset. Cells did not show similarity by batch by visualizing clustering or in downstream analyses such as differential expression analysis. Batch effects corresponding to library preparation and sequencing runs were evaluated by visual inspection of UMAP embeddings and were not associated with genotype, age, or region. Attempts to eliminate batch effect were made using Harmony (v1.2.3)^46^ but resulted in over-smoothing of variation within cell types and resulted in no differences being detected across groups. It was ultimately decided to not employ batch correction as gene expression trends do not segregate by batch.

### Identification of cell type-specific DEGs

Differential expression analyses were performed for each cell type at every timepoint between WT and transgenic for each spinal region using the FindMarkers command in Seurat. The default Wilcoxon Rank Sum test was used for pairwise comparisons. Differentially expressed genes were filtered for those with at least log fold change of 1 and less than 0.05 adjusted p-value. Identification of convergent DEG trajectory was performed by listing all the DEGs for each comparison and plotting their expression over time using pHeatmap (v1.0.12)^47^. Cell expression patterns for each cell type were hierarchically clustered in pHeatmap. Clusters of cell expression patterns were identified by calculating optimal silhouette based on the maximum silhouette width per number of clusters k and cutting the tree at the optimal k value. Clusters of cell expression identified for each cell type are then compared for similarity with other cell types visually. Module scores were calculated for genes in each cell type cluster and reclustered in pHeatmap to identify the eight major shared DEG expression patterns across cell types.

### Identification of cell type specific aging gene networks

Aging gene co-expression networks were constructed for each cell type and genotype using the hdWGCNA (v0.4.4) R package^23^. The Seurat object containing postnatal snRNA-seq data was filtered to exclude embryonic neurons, iPSCs, and radial glia. Cells were split by both cell type and genotype into sub-objects using the SplitObject() function. For each object, the top 4,000 most variable genes were identified using Seurat’s FindVariableFeatures(), and data were normalized, scaled, and subjected to PCA and UMAP for visualization and downstream grouping. Metacells were constructed using the MetacellsByGroups() function, with cells grouped by age and spinal region (age_spinal_region). Metacells were constructed in UMAP space using k = 3 nearest neighbors and allowing up to 10 shared cells between metacells. Normalized expression values were then aggregated across metacells. Soft-thresholding powers were tested using TestSoftPowers(), and scale-free topology fit curves were plotted. For each cell type/genotype combination, the soft-thresholding power was selected based on the scale-free topology criterion. One cell type (Motorneuron_SD) was removed from network construction due to low quality or non-convergence. Gene co-expression networks were constructed with ConstructNetwork(), without re-running SetDatExpr() since it was already performed.

Dendrograms of hierarchical clustering of genes were generated for all modules. Module eigengenes (MEs) were computed across all metacells using ModuleEigengenes(), grouped by age_spinal_region. Both harmonized and raw MEs were extracted using GetMEs(). To evaluate gene membership within modules, eigengene-based connectivity (kME) was calculated using ModuleConnectivity(). Only modules with eigengenes showing Pearson correlation coefficients ≥ |0.3| with age were considered aging-associated modules. These aging modules were used for downstream transcriptional regulator analysis and functional enrichment.

### Microglia aging trajectory analysis

Microglia were subset from the integrated dataset based on cell type annotation (“Microglia”) generated during initial preprocessing and clustering. No additional quality control filtering was performed at this stage.

The subsetted cells were reclustered using Seurat following standard workflows, including data scaling, principal component analysis (PCA), and shared nearest neighbor graph-based clustering. Trajectory inference was performed using Monocle3 v1.3.1^48–51^. The Seurat object was converted to a Monocle 3 *cell_data_set* object, and dimensionality reduction was performed using UMAP. Cells were ordered in pseudotime using the *learn_graph* and *order_cells* functions. The root node was defined as the node farthest within the cluster enriched for the earliest timepoints (P25–P50), such that pseudotime reflects progression from early to late aging states. Default parameters were used unless otherwise specified.

### Gene ontology

Gene Ontology (GO) enrichment analysis was performed using the WebGestaltR package v0.4.4^52^ in R. Overrepresentation analysis (ORA) was used to identify significantly enriched Gene Ontology terms in the Biological Process (BP), Cellular Component (CC), and Molecular Function (MF) categories. For each gene list of interest, GO enrichment was performed using WebGestaltR with the following parameters:Organism - Mus musculus, Gene type - gene symbols, Reference set - entire mouse genome, Minimum and maximum gene set size - 5 and 2,000, respectively, Significance method - False Discovery Rate (FDR), adjusted using the Benjamini-Hochberg (BH) method, Significance threshold - FDR < 0.05, Redundant term reduction - setCoverNum = 10 was used to reduce redundancy among enriched GO terms.

### Regulon construction custom scoring

Jaccard indices were calculated in R by dividing the intersections by the union of the two gene sets. Results are presented as a heatmap using the pHeatmap package. Transcription factor regulons were inferred for each cell type using the pySCENIC pipeline (v0.11.2)^53^ in Python. Gene expression matrices were generated from metacell-aggregated expression data derived from hdWGCNA-preprocessed Seurat objects. Only genes present in both the expression matrix and the motif databases were retained for analysis. For each aging gene module of each cell type, gene regulatory networks (GRNs) were first inferred using the GRNBoost2 algorithm implemented in the arboreto package, using a curated list of mouse transcription factors (mm_mgi_tfs.txt). This generated a ranked list of TF–target gene relationships (adjacencies). Putative modules were extracted from the adjacencies using pySCENIC’s modules_from_adjacencies() function. These modules were then pruned using motif enrichment analysis with the cisTarget motif databases (v9, mm10), which include genome-wide motif rankings in .feather format. Motif annotation was performed using the motifs-v9-nr.mgi-m0.001-o0.0.tbl annotation file. The pruned network was converted into a list of regulons—TFs with their target gene sets supported by motif evidence. Regulon activity was quantified using AUCell, which computes the area under the recovery curve (AUC) for each regulon across all metacells. AUC matrices were saved and visualized as heatmaps using Seaborn’s clustermap() function. Optionally, AUCell matrices were binarized using a custom binarize_matrix() function, and binarized heatmaps were generated to visualize regulon ON/OFF states across metacells. All pySCENIC analyses were performed separately for each cell type using 8 CPU threads per run. Intermediate and final outputs including adjacencies, modules, regulons, AUC matrices, and plots were saved per cell type in structured output directories for downstream integration with aging-related gene modules. To further assess the activity of transcription factor regulons identified using pySCENIC, we re-scored regulon activity using the AUCell algorithm in R (AUCell v1.18.1). Gene expression matrices derived from metacell-aggregated data were loaded for each cell type, and pySCENIC-inferred regulons were parsed from CSV files containing transcription factor–target gene relationships. Regulons were represented as lists of target gene sets for each transcription factor. For each metacell expression matrix, genes were ranked within each cell, and AUCell_buildRankings() was used to compute ranked gene lists.

AUCell scores were calculated for all regulons using the AUCell_calcAUC() function, with the aucMaxRank set to the top 5% of ranked genes. The resulting regulon-by-cell AUCell score matrices (continuous activity scores) were saved and visualized using the pHeatmap package. To facilitate binary classification of regulon activity, a threshold was computed for each regulon using the 95th percentile of its AUCell distribution across all metacells. Regulons were considered active in a cell if the AUCell score exceeded this threshold. These binary activity matrices were also saved and visualized as heatmaps. Cell metadata (age and spinal region) were extracted from column names in the expression matrices. This metadata was used to annotate heatmaps and to sort cells by age. Heatmaps of both continuous and binarized AUCell scores were generated, with and without Z-score scaling across cells. Rows (regulons) were hierarchically clustered, while columns (cells) were ordered by age. Transcription factor scores are computed by the sum of weighted values. These values include the enrichment of target genes of the transcription factors in the aging gene module (weight = 0.8), the correlation of the AUC to eigengene expression (weight = 0.3), the correlation of the transcription factor expression to the module eigengene (weight = 0.3), and the magnitude of the expression level of the transcription factor (weight = 0.6).

### RNAscope In Situ Hybridization

Mice were perfused with PBS until organs appeared pale, followed by 4% paraformaldehyde until stiff. Spinal cords were dissected and post-fixed at 4 °C overnight, then washed three times with PBS the following day. Tissues were stored in 30% sucrose with 0.1% sodium azide at 4 °C until sectioning. Major spinal regions were identified and dissected as described for nuclei isolation. Spinal regions were embedded in optimal cutting temperature (OCT) compound and frozen. After temperature equilibration in a Leica CM3050S cryostat, tissues were mounted, trimmed, and sectioned at 14 µm thickness. Sections were stored in antifreeze solution prepared by dissolving 85.6 g sucrose and 0.66 g MgCl₂ in 500 mL of 1× PBS and adjusting the final volume to 1 L with glycerol. Sections were mounted on charged glass slides, and in situ hybridization was performed using the RNAscope™ Multiplex Fluorescent Detection Kit v2 (ACDbio, Cat# 323110) according to the manufacturer’s instructions. Human SOD1 mRNA was detected using the RNAscope™ 2.5 VS Probe Hs-SOD1 (ACDbio, Cat# 1147999-C1). Nuclei were stained with VECTASHIELD Vibrance Antifade Mounting Medium containing DAPI (Vector Laboratories, Cat# H-1800-2) and imaged using an EVOS M5000 fluorescence microscope.

### Genomic and Mitochondria DNA

Mouse spinal cord regions were homogenized using plastic tissue crusher in 500 ul of lysis buffer (25 mM Tris-HCl (pH 7.8), 10 mM NaCl, 0.2 mM EDTA, 0.0125% SDS, 200 ug/ml Proteinase K (New England Biolabs, Cat# P8107S)). Samples were incubated overnight at 55 °C with 300 rpm in a tube shaker. RNAse A (New England Biolabs Cat#T3018L) was added for a final concentration of 20 ug/ml for 15 minutes at 37 °C after lysis. 500 ul of 1:1 phenol-chloroform was added to the lysate and vortexed for 15 seconds, then centrifuged at 12,000 g for 15 minutes. The aqueous phase was extracted and added to the same amount of isopropanol. Sodium acetate (pH 5.2) was added to a final concentration of 0.3 M. The sample was mixed by inverting then incubated at -20 °C for 30 minutes. The samples were centrifuged at 12,000 g for 10 minutes at 4 °C. The supernatant was then discarded and the pellet washed with 70% ethanol. DNA pellets were air dried for 10 minutes at room temperature then eluted in water for 10 minutes at 37 °C. DNA concentrations were measured using Nanodrop and equalized to 50 ng/ul by adding water. 2 ng of total DNA were amplified by qPCR using primers for mitochondrial and nuclear DNA to determine mitochondrial to nuclear DNA ratio. *Mt-Nd1* and *mt-Nd6* were averaged to assess total mitochondria DNA. *B2m* and *Tert* were averaged to represent total nuclear DNA. Delt CT (average mitochondria – average nuclear) was calculated to represent the ratio between mitochondria and nuclear DNA. One biological replicate of the control condition was used to calculate delta delta CT (ddCT). Fold change was calculated by taking the 2^-ddCT^.

### qPCR

Flash frozen spinal cords were dissected for spinal region as described in the nuclei isolation section. Spinal cord sections were dounced in 1 ml of Trizol until homogenized. RNA isolation procedures were performed using Qiagen RNeasy Kit (Qiagen Cat#74104). cDNA synthesis was performed using NEB Protoscript II First Strand cDNA Synthesis Kit (New England Biolabs, Cat# E6560L). qPCR was performed using SYBR Green chemistry (Applied Biosciences, Cat# 4309155) in a Bio-Rad CFX384 Real Time PCR system. Fold changes were analyzed in Microsoft Excel. For RNA expression, mouse *U6* was used as housekeeping gene as this gene remains at constant levels during aging and ALS progression.

### Western Blot

Spinal cord tissues were weighed and RIPA buffer containing protease inhibitor cocktail (Roche) were added to a total of 100 µg tissue per ul of buffer. Tissues were homogenized by sonication. Homogenates were run on 10% polyacrylamide gel with Laemmli (Bio-Rad, Cat# 1610737EDU) +5% betamercaptoethanol at a 1:1 ratio. Gels were transferred to nitrocellulose membrane (Bio-Rad Cat#1704158) using Bio-Rad Trans-Bot Turbo Transfer system. Proteins were quantified using Ponceau staining. Li-cor Intercept blocking buffer was used to block the membranes. All antibodies used were diluted at a 1:1000 ratio in TBS-T (Tris-buffered saline (0.1% Tween)). Primary antibody incubation was done overnight at 4 °C with gentle shaking (SOD1 Antibody, Cell Signaling, Cat#2770, GAPDH-peroxidase, Millipore Sigma, G9295). Membranes were washed three times with TBS-T with 5-minute incubations. HRP-conjugated secondary antibodies (Anti-mouse IgG HRP-linked antibody, Cell Signaling, Cat# 7076P2, Anti-rabbit IgG HRP-linked antibody, Cell Signaling, Cat# 7074P2) were diluted 1:10000 in TBS-T and incubated for two hours. Then washed three times with TBS-T with 5-minute incubations between each wash. Blots were developed using Bio-Rad Clarity Western ECL substrate (Bio-Rad, Cat#1705061) and imaged on Bio-Rad Chemidoc Chemiluminescence imager. Signal quantification was analyzed in Fiji using the gel analyzer plugin. An empty area was selected as background. The same rectangular ROI area was used to measure each band. Band intensities were quantified by measuring integrated density values using a fixed rectangular ROI. The background AUC was subtracted from the band AUC measurements. Measurements were divided from Gapdh measurement or from whole lane Ponceau AUC.

### Statistical Testing

Statistical analyses of qPCR and western blot data were performed in R (version 4.2.2). Fold changes were log₂-transformed to normalize the distribution. Unpaired two-tailed t-tests were used to compare groups. p-values were adjusted for multiple comparisons using the Benjamini-Hochberg method. Data are presented as mean ± standard deviation, and corrected p-values < 0.05 were considered statistically significant. Three biological replicates were used per group.

---

**Supplementary Figure 1.**
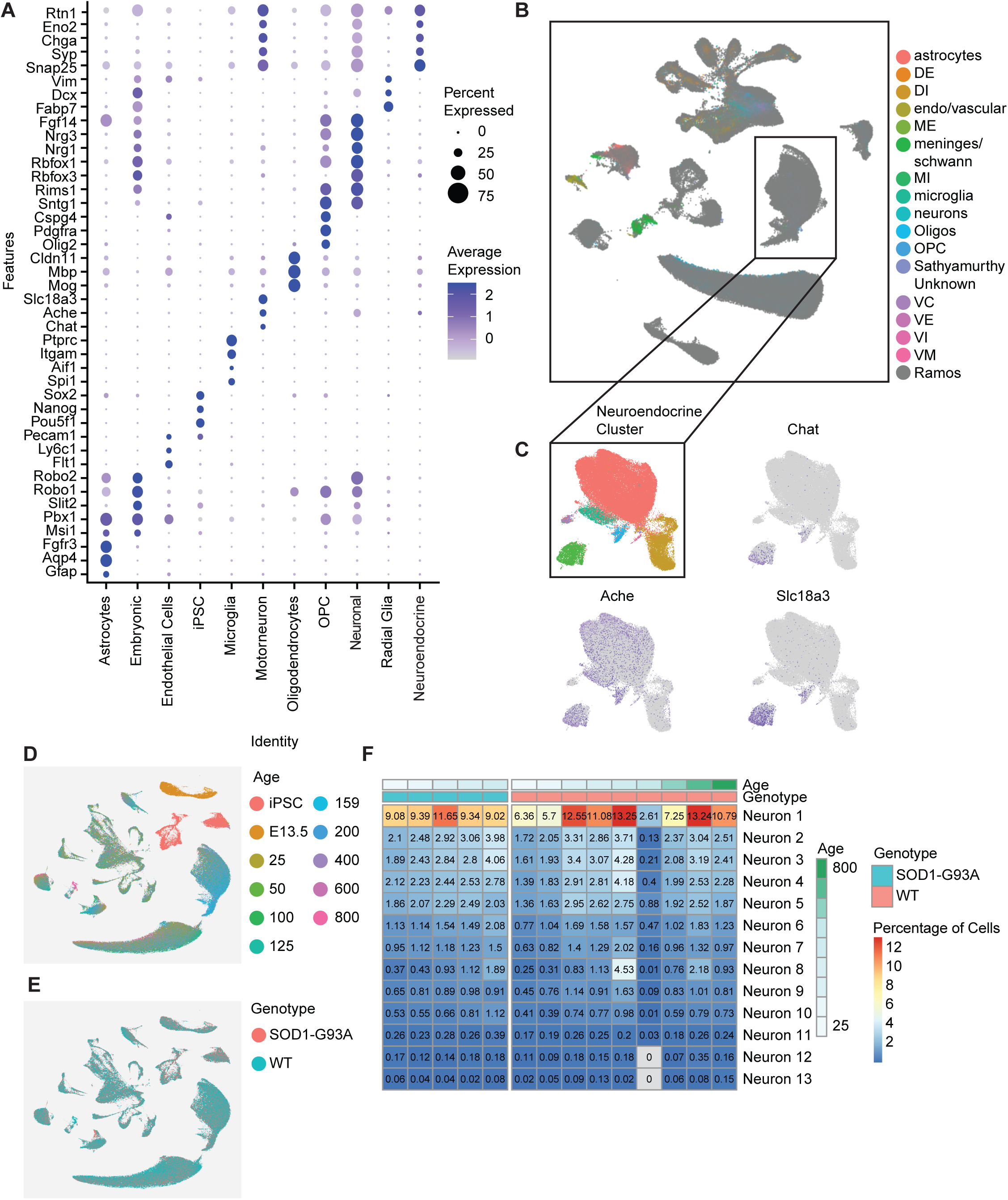
**Cell type annotation and validation across lifespan and genotype.** a, Dot plot of canonical marker genes used to annotate major spinal cord cell types, showing specificity and relative expression across clusters. b, Integrated UMAP embedding harmonized with annotations from Sathyamurthy et al. (2018), demonstrating concordance of major neuronal and glial identities across datasets. c, Reclustering of neuroendocrine populations to resolve motor neurons, identified by expression of cholinergic markers (*Ache*, *Chat*, *Slc18a3*). UMAP of transcriptomic atlas colored by age (d), and genotype (e). f, Heatmap of neuronal subtype proportions across ages in WT and SOD1-G93A mice.

**Supplementary Figure 2.**
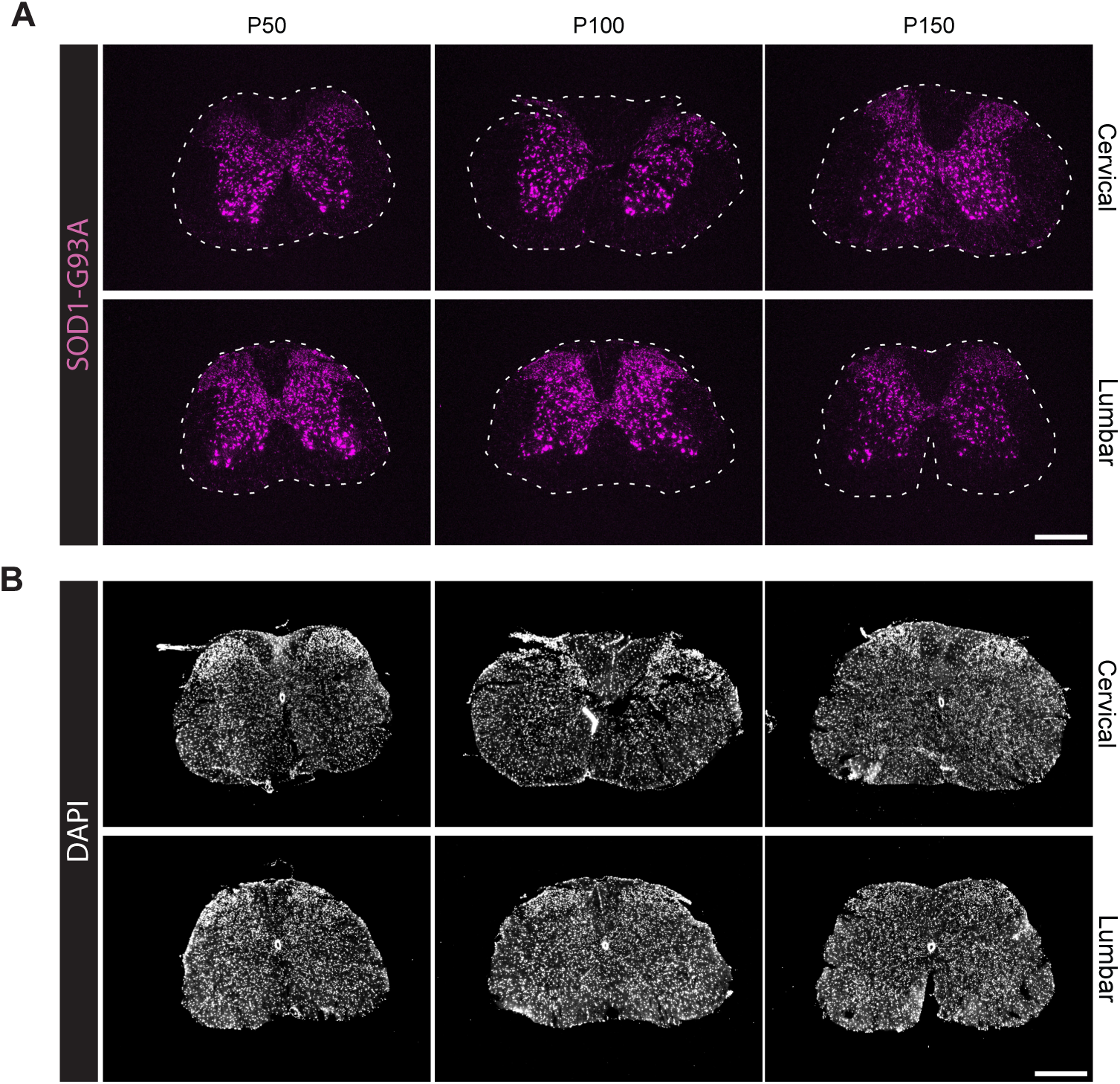
**Spatial SOD1-G93A expression across cervical and lumbar spinal sections.** a, RNAscope detection of *SOD1-G93A* transcripts in cervical (top) and lumbar (bottom) spinal cord at presymptomatic (P50), onset (P100), and symptomatic (P150) stages, showing spatially distributed expression across grey matter. Scale bars, 500 μm. b, Nuclear counterstaining of DAPI corresponding to panel a, confirming predominantly grey matter cell localization of *SOD1-G93A* transcripts. Scale bars, 500 μm.

**Supplementary Figure 3.**
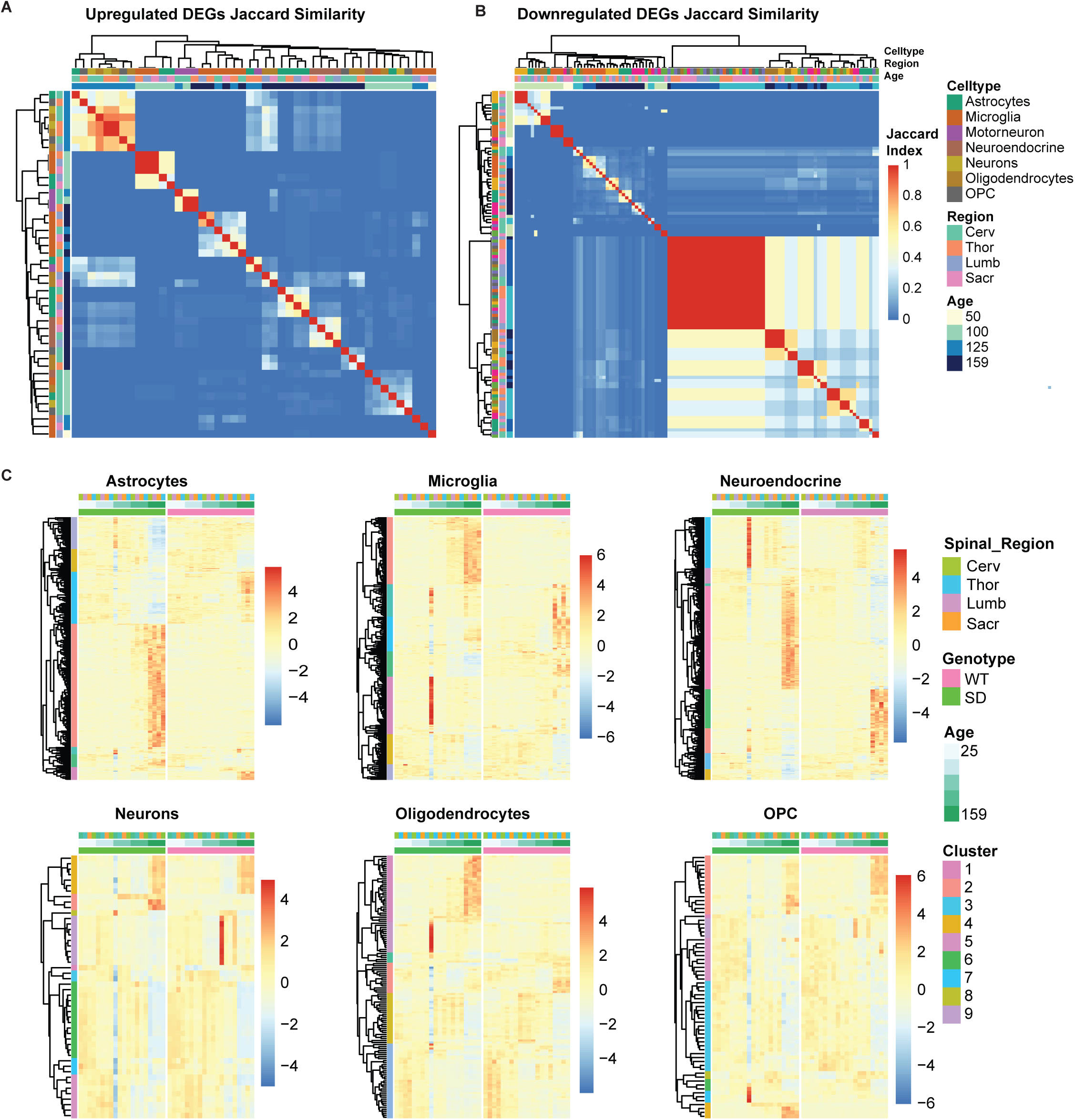
**Cross-cell type similarity and clustering of differential expression programs.** Jaccard similarity of upregulated (a) and downregulated (b) DEGs across cell types at each disease stage, showing shared transcriptional responses. c, Hierarchical clustering of DEGs in WT (left) and SOD1-G93A (right) across major cell types, revealing shared temporal gene expression patterns.

**Supplementary Figure 4.**
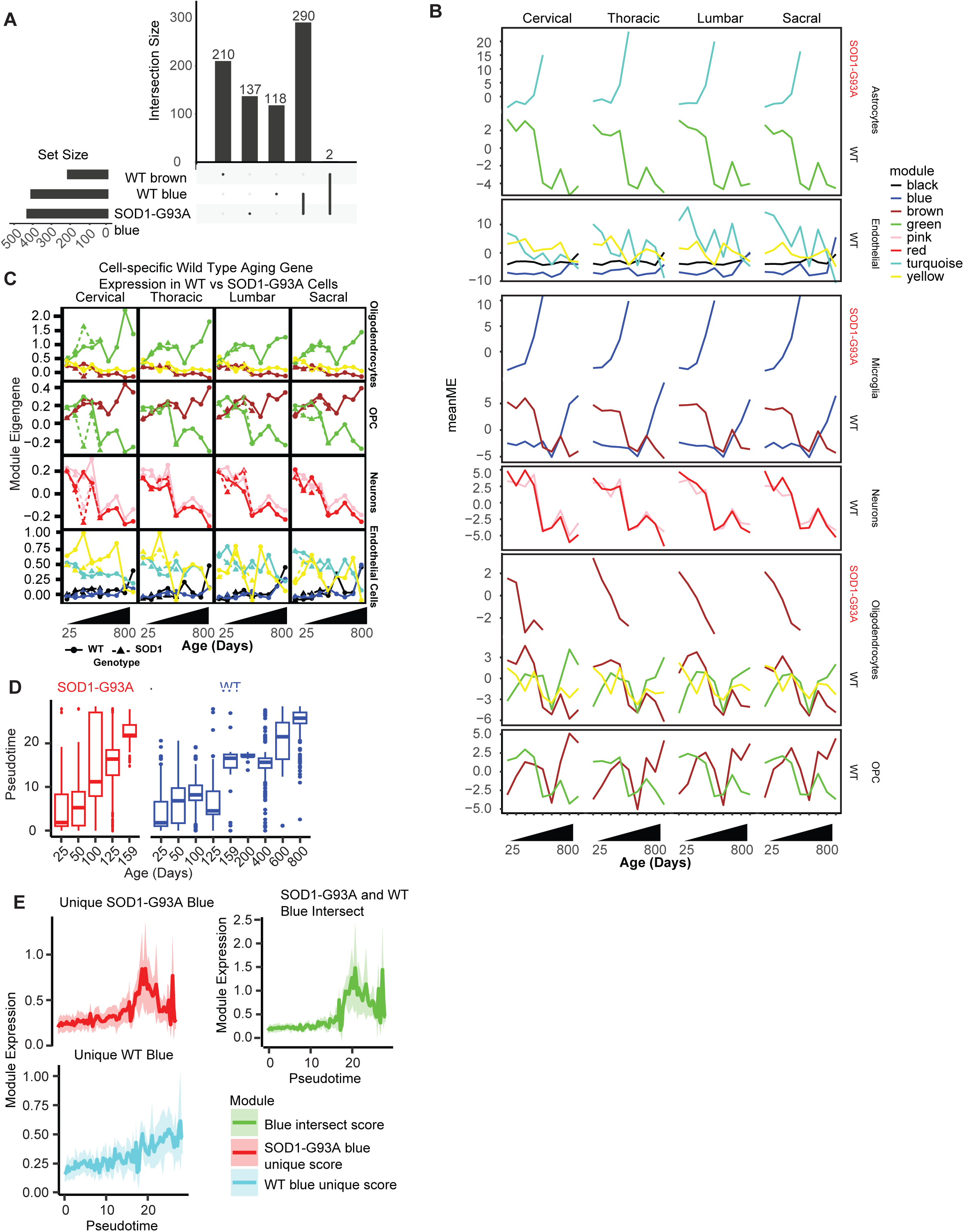
**Preservation of WT aging programs in SOD1-G93A cells.** a, UpSet plot of gene overlap between WT and SOD1-G93A microglial aging modules, showing predominant overlap of their respective blue modules. b, Expression of aging-associated module eigengenes across time in four spinal cord regions, showing regionally consistent but cell type–specific aging trajectories. c, Temporal trajectories of WT aging modules in WT (solid) and SOD1-G93A (dashed) oligodendrocytes, OPCs, neurons, and endothelial cells, demonstrating overall preservation of normative aging dynamics. d, Microglial pseudotime trajectories across the lifespan, showing accelerated progression in SOD1-G93A cells relative to chronologically age-matched WT cells. e, Expression of shared and genotype-specific microglial aging module genes across pseudotime.

**Supplementary Figure 5.**
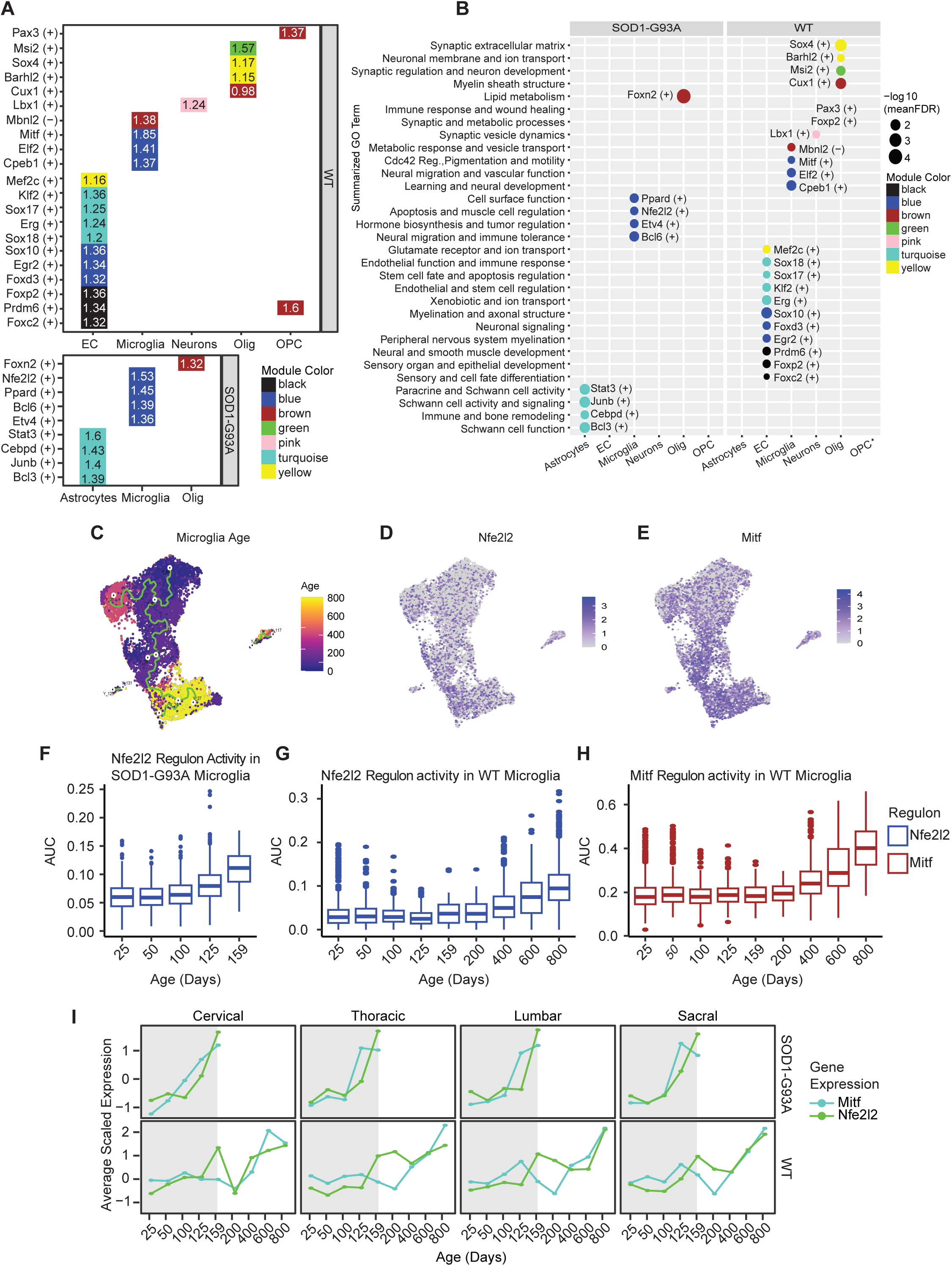
**SOD1-G93A disrupts MITF and Nrf2 regulatory programs in microglia.** a, Top-scoring aging-associated regulons for each cell type in WT (top) and SOD1-G93A (SD, bottom), ranked by weighted regulon scores per cell type and genotype. (+) / (-) refers to directionality of regulon correlation to aging module eigengene. b, Gene Ontology enrichment of top regulons per cell type and genotype. c, UMAP of microglia colored chronological age. Green line denotes pseudotime axis and branch points between P25 and 800. d,e, Feature plots of *Nfe2l2* and *Mitf* expression in microglia, showing increased expression correlated with age. f–h, Temporal pattern of Mitf and Nrf2 regulon activity in WT and SOD1-G93A microglia, illustrating gradual increase with WT aging and disease progression. i, Regional and temporal expression of *Mitf* and *Nfe2l2* across the rostrocaudal axis, showing similar expression pattens across regions, aging, and accelerated induction in SOD1-G93A cells.

## Supplementary Tables

Supplementary Table 1. Cell type–resolved differential expression analysis across stages and genotypes.

Supplementary Table 2. Gene membership of temporally defined DEG clusters for each cell type.

Supplementary Table 3. Gene Ontology enrichment for DEG clusters across cell types and disease stages

Supplementary Table 4. Cell type–specific aging module gene sets derived from co-expression analysis.

Supplementary Table 5. Gene Ontology enrichment of cell type–specific aging modules.

Supplementary Table 6. Gene Ontology enrichment of transcription factor regulon targets associated with aging programs.

